# Elucidating the genome of *Emydura subglobosa* through manual annotation and computational biology tools

**DOI:** 10.1101/2021.06.24.449536

**Authors:** Megan R. Yu

## Abstract

Rapid advancements in automated genomic technologies have uncovered many unique findings about the turtle genome and its associated features including olfactory gene expansions and duplications of toll-like receptors. However, automated technologies often result in a high frequency of errors through the process of assembly and annotation and highlight the need for manual annotation. In this study, we have manually annotated four genes of the red-bellied short-neck turtle (*Emydura subglobosa*), an understudied outgroup of turtles representing a diverse lineage. We improved upon initial *ab initio* gene predictions through homology-based evidence and generated refined consensus models. Through functional, localization, and structural analyses of the predicted proteins, we have discovered conserved genes encoding proteins that play a role in C21-steroid hormone biosynthetic processes, Vitamin A uptake, collagen/elastin integrity, tumor suppression, and fatty acid catabolism. Overall, these findings further our knowledge about the genetic features underlying turtle physiology, morphology, and longevity, which could have important implications for the treatment of human diseases and evolutionary studies.

## Introduction

Turtles have been on Earth for over 210 million years, with the earliest estimated stem-turtle originating in South Africa approximately 260 million years ago [1]. Compared to other eukaryotic species, turtles have minimal diversity (360 living species) and over 50% are threatened with extinction [2]. Understanding turtle biodiversity is difficult because they have low mitochondrial deoxyribonucleic acid (mtDNA) variation and low evolutionary change on a molecular level, which could be due to their slow maturity and unconstrained hybridization [3, 4]. With that in mind, advancements in turtle genomics have only begun since 2013, despite evidence showing that turtles have been around for over 210 million years.

The field of turtle genomics demonstrated major breakthroughs beginning in 2013 with the first three turtle genomes and continues to progress to this day. In 2013, the entire genome of the western painted turtle (*Chrysemys picta bellii*) was sequenced, assembled, and examined through comparative genomic and phylogenetic analyses, highlighting that turtles are sister to archosaurs and evolve very slowly at the sequence level. The painted turtle possessed genes that allowed them to tolerate reduced oxygen concentrations and freezing temperatures; analyses showed convergent evolution of tooth loss based on pseudogenization of those specific genes [5]. Furthermore, in this same year, the soft-shell turtle (*Pelodiscus sinensis*) and green sea turtle (*Chelonia mydas*) genomes were sequenced, thereby exhibiting a large expansion of olfactory genes to enhance their sense of smell [6]. The Pinta Island tortoise (*Chelonoidis abingdonii*) and Aldabra giant tortoise (*Aldabrachelys gigantea*) were sequenced in 2019 and provided insight into longevity and age-related diseases, specifically through duplications in age-related cell repair and cancer resistance genes. These tortoises also had favorable mutations in DNA repair genes that carried through evolution [7]. Finally, whole duplications of toll-like receptor (TLR) genes were shown to trigger downstream immune response in the desert tortoise [8]. With the advent of new, high throughput sequencing technologies and annotation tools, we have begun to uncover many discoveries about turtles.

In this paper, manual gene annotation will be performed on genes of the red-bellied short-neck turtle (*Emydura subglobosa*), a diverse species whose genome has not been fully annotated. Genome sequencing, genome assembly, and automated annotations have been conducted on *E. subglobosa*; however, there has been a lack of manual genome annotation which could unravel important evolutionary information and molecular mysteries encoded in its genes. This project aims to improve upon *ab initio* predictions and annotations provided by automatic annotation tools, such as AUGUSTUS and Apollo, and conduct both structural and functional analysis [9, 10]. While genomic annotation involves several types of data and methods, our strategy is exclusively based on homology and computational biology, using tools such as Basic Local Alignment Sequence Tool (BLAST), Constraint-based Multiple Alignment Tool (COBALT), InterPro, WoLF PSORT (Protein Subcellular Localization Prediction), Transmembrane Helices Hidden Markov Models (TMHMM), SWISS-MODEL, and STRING. Precise elucidation of the features encoded in the *E. subglobosa* genome gives insight into both the structure and function of these genes and their relationship to other eukaryotic species across evolutionary time, which can further advance conservation efforts and clinical medicine therapies.

## Methods

### Genome Sequencing and Assembly

We utilized a prebuilt genome that was previously sequenced, assembled, and published on the NCBI Assembly database by the name of Emydura_subglobosa-1.0 [11, 12]. Under the McDonnell Genome Institute at the Washington University School of Medicine, a sample of blood from a male, red-bellied short-neck turtle (*E. subglobosa*) at the St. Louis Zoo in St. Louis, MO was completely sequenced using 10X Genomics Chromium technology [11]. This high throughput sequencing technique uses single-cell RNA-sequencing (scRNA-seq) by capturing single cells on a gel micro-bead in emulsion labeled with oligonucleotides complementary to adaptors [13]. The fully sequenced Emydura_subglobosa-1.0 genome was approximately 2.6 Gb long and had 61x coverage. The genome was then assembled at the scaffold level using Supernova v. 2.0.1 [11]. This method is a whole genome *de novo* assembly process that utilizes the library created by Chromium to create large contigs as reads are assembled together [14]. The full Emydura_subglobosa-1.0 genome—43,399 scaffolds and 56,575 contigs long—contained a 351 Kb N50 contig and 44.8 Mb N50 scaffold which could be used for further structural and functional analysis [11].

### *Ab initio* Gene Prediction from AUGUSTUS

In this study, we used the AUGUSTUS gene prediction as a starting point for structural and functional analysis of the *E. subglobosa* genome. AUGUSTUS is based on the Hidden Markov Model (HMM) and performs *ab initio* gene prediction to define structural elements such as coding and non-coding regions, splice sites, and intergenic length distributions [9]. The AUGUSTUS prediction was initially evaluated through homology-based evidence and subsequently edited and/or validated through genomic biology tools.

### Local Fragment Alignments through BLASTP

NCBI Protein Basic Local Alignment Sequence Tool (Protein BLAST or BLASTP) was used to compute alignments of the predicted query sequence against a protein database and calculate statistical significance of alignments based on similarities. This tool allowed us to search for optimal alignment fragments with other species based on the maximal segment pair (MSP) and the pairwise alignment algorithm [15]. After selecting a gene of interest on Apollo, we inputted the FASTA sequence of the predicted gene into the query search of standard Protein BLAST. The default settings of non-redundant protein sequences (nr) database and blastp algorithm were used. From BLAST, we were able to obtain important outputs including the description of top-hit homologs that shared common sub-sequences or fragments, percent identity for exact residue matches, query coverage, maximum high-scoring segment pair (HSP) score with no gaps, and E value showing statistical probability of chance HSP alignments [16]. Selection of the same isoform across the top 7 to 10 highest-matching species was done to avoid duplicating the same alignments. Only E values less than 0.1 and near zero were considered to ensure the low probability of the BLAST alignments occurring due to chance. This gave us increased confidence that the alignment with the homolog was real, so we could deduce that the two species evolved from a common ancestor. We also considered genes with a high percent identity and query coverage greater than 70%, so that we could find high conservation across different species. After selecting the top 7 to 10 best subject hits (same isoform) based on the pairwise alignment, we downloaded the complete FASTA sequence, rather than the aligned FASTA sequence, to obtain the entire sequences of all subjects in case the predicted model missed any information outside the aligned fragment. We pasted the predicted AUGUSTUS sequence to the top of the seqdump.txt file. Through these considerations, we could be confident about the local alignment results from Protein BLAST and move on to further analysis.

### Multiple Sequence Alignment using COBALT

While Protein BLAST performs local alignment with no gaps, this algorithm does not guarantee optimal alignments due to potential evolutionary insertions and deletions (gaps). Therefore, Constraint-based Multiple Alignment Tool (COBALT) was used to detect other highly conserved, homologous protein sequences, provide insight into protein function, structure, and evolution, and identify sequencing errors and regions to be edited. Based on the cluster alignment (CLUSTAL) algorithm, COBALT uses pairwise alignment to calculate distance matrices, which are then continuously aligned based on a guide tree [17]. We uploaded the seqdump.txt FASTA file to COBALT using default settings. From the COBALT output, we were able to visualize the structural schematic of the highly conserved (red) and variable regions (gray) of the protein sequence. At the bottom, we were able to see the exact protein sequences across conserved homologs and figure out which sequences were added, missing, or different in the query compared to the other validated sequences.

### Evidence-based Genome Editing and Annotation using Apollo

If the query showed distinct differences compared to the conserved protein homologs in other species, genome editing in Apollo was used to refine the AUGUSTUS prediction and generate a consensus model [10]. All edits were made from the perspective of the query compared to the homologous proteins. Gaps between exons or long gray exons were evidence of missing sequences or extra sequences in the coding region of the predicted gene model, respectively. For these missing sequences, we used the Search tab, pasted the amino acid sequence that was present in the other homologs or found in the non-coding region nearby, selected “Blat protein,” and searched. If a match was found, we made a new exon encompassing that specific amino acid sequence so that it was no longer missing. For the long gray regions representing low conservation that were not present in other homologs, these extra exons were removed from the predicted protein by either deleting the entire exon or shortening it. If the predicted model was shorter than the other conserved sequences, this was evidence that the gene model was truncated and needed to be extended. New exons were created by 1) extending current exons, 2) splitting the current exon and dragging the new one to the appropriate region, or 3) merging two different genes. For short missing or extra sequences found in the middle of an exon, these were left alone since it could be due to evolutionary insertions or deletions. All edits were carefully completed to maintain the open reading frame for proper transcription. The gene model was iteratively refined through the use of the three tools—BLAST, COBALT, and Apollo—to generate a new consensus model that was comparable to the conserved genomes based on homology.

### Functional Analysis

To determine the function(s) of the protein, we used InterPro, AmiGO 2, and STRING v. 11.0. The purpose of InterPro was to determine protein function based on similar sequence motifs and domains [18]. Because domains are functional parts of a protein and are highly conserved, inferring function through domain searches allowed us to classify the protein of interest into families. We pasted the FASTA sequence of the consensus model into the “Search by sequence” box and used default settings. The protein family membership was noted. We visualized the different families, domains, superfamilies, and predictions that stretched along the length of the gene at varying lengths. Gene ontology (GO) terms for biological processes, molecular functions, and cellular components were also outputted at the domain level and recorded. In addition, AmiGO 2 was used to complement InterPro’s GO terms by generating top hits of GO terms associated with the protein itself, rather than at the domain level [19]. We searched the homologous gene name associated with the protein and evaluated the ontology graph under the Graph Views tab, which gave insight into the larger biological process that the protein was involved in.

For further functional analysis, we used STRING v. 11.0 to predict direct and indirect protein-protein interactions [20]. Under the “Protein by sequence” sidebar, the sequence was pasted into the “Amino Acid Sequence” box and the organism was “*Homo sapiens*.” Humans were used as the organism in order to look into conservation of interacting proteins and see if these interaction networks were conserved in turtles. The outputted network from STRING highlighted several key interactions and functional partners. Under the “Viewers” tab, we examined the gene cooccurrence: dark red squares corresponding to gene families whose occurrence patterns showed strong homology across species, light red squares corresponding to gene families whose occurrence patterns showed weak homology across species, and missing squares corresponding to gene families whose occurrence patterns showed no direct homologies across species.

### Subcellular Localization

To identify the subcellular localization, different complementary tools were utilized, including WoLF PSORT (Protein Subcellular Localization Prediction), Transmembrane Helices Hidden Markov Models (TMHMM) Server v. 2.0, SignalP-5.0 Server, Phobius, and TargetP-2.0 Server. WoLF PSORT is a tool that predicts protein localization sites based on the primary amino acid composition [21]. We selected the “Animal” organism type, kept “From Text Area” as default, and pasted the FASTA sequence of the consensus model into WoLF PSORT. We noted the highest scoring matches based on percent identity which predicted where the protein was localized. TMHMM Server v. 2.0 further complemented results from InterPro and WoLF PSORT by prediction of transmembrane helices (TMHs). TMHMM uses the probabilistic Hidden Markov Model (HMM) developed from observed sequences of proteins with known functions [22]. We pasted the FASTA sequence using default settings. The top schematic summarized discrete regions within the protein. Probabilities greater than 0.75 were considered significant in indicating segments of the protein that lie inside, outside, or within the membrane. Additionally, SignalP-5.0 Server was used to predict the signal peptide cleavage site that targets the protein to its correct location [23]. We pasted the FASTA sequence of the protein and used the default settings of “Eukarya” organism group and “long output” format. Probabilities greater than 0.5 suggested that the gene encodes a signal peptide; probabilities greater than 0.75 gave us greater confidence regarding the presence of a cleavage site. To confirm the outputs of TMHMM and SignalP, we used Phobius to predict TMHs and signal peptides [24]. Again, we pasted the FASTA sequence with default settings, and probabilities greater than 0.75 were considered significant. Finally, TargetP-2.0 was used to identify N-terminal presequences, signal peptides, and transit/targeting peptides [25]. We pasted the FASTA sequence with default settings of “Non-Plant” organism group and “long output” format. Probabilities greater than 0.5 suggested that the gene encodes an N-terminal presequence, signal peptide, or transit peptide; probabilities greater than 0.75 gave us greater confidence. Overall, these tools gave insight into where the protein localizes within the cell.

### Protein Modeling and Structure

To complement the TMHMM results, we also used SWISS-MODEL to generate a three-dimensional (3D) protein structure based on homology [26]. We pasted the FASTA sequence into the “Target Sequence” and built the model. From the SWISS-MODEL output, we were able to obtain information regarding oligo-state, ligands, global (GMQE) and local (LMQE) model quality estimations, model-template alignments corresponding to secondary structures, comparison with non-redundant set of protein database (PDB) structures, and an overall 3D protein structure model. We examined the model with the GMQE closest to 1 and confirmed results with TMHMM regarding transmembrane helices.

To validate the model, we examined the LMQE, comparison with non-redundant set of PDB structures, Ramachandran Plot, and MolProbity results. The LMQE shows pair residue estimates, and scores <0.6 were considered low-quality. For the graphical comparison with non-redundant set of PDB structures, the dark gray represented the QMEAN score for experimental structures of similar size; we considered a good model to fall within the gray range. The Ramachandran Plot under “Structure Assessment” shows the probability of a residue having a specific orientation. We validated the model if it had a Ramachandran Favoured value close to 100%, Ramachandran Outliers value close to 0%, and MolProbity Score close to 0. The final tool we used to validate the homology structure was PSIPRED 4.0, in which we pasted the protein sequence and submitted using default settings. PSIPRED predicts secondary structure, which also complemented TMHMM results [27].

### Phylogenetic Tree Analysis

We used the phylogenetic tree functions of BLAST and COBALT to scrutinize the evolution of the protein across various species and see the graphical relation between genomic sequences in tree format. BLAST and COBALT use pairwise alignment and multiple sequence alignment, respectively, to construct the tree [15, 17]. More similar sequences branched closer to the right, and less similar sequences branched closer to the left. We built the phylogenetic tree and expanded out organism sets by including disparate species into the seqdump.txt file with lower percent identities. This was to confirm the relative similarities between homologs versus distant species. Through BLAST and COBALT, we were thus able to determine how similar the other homologs aligned to the predicted sequence. Table 1 lists all the tools used in the methods.

**Table 1.**
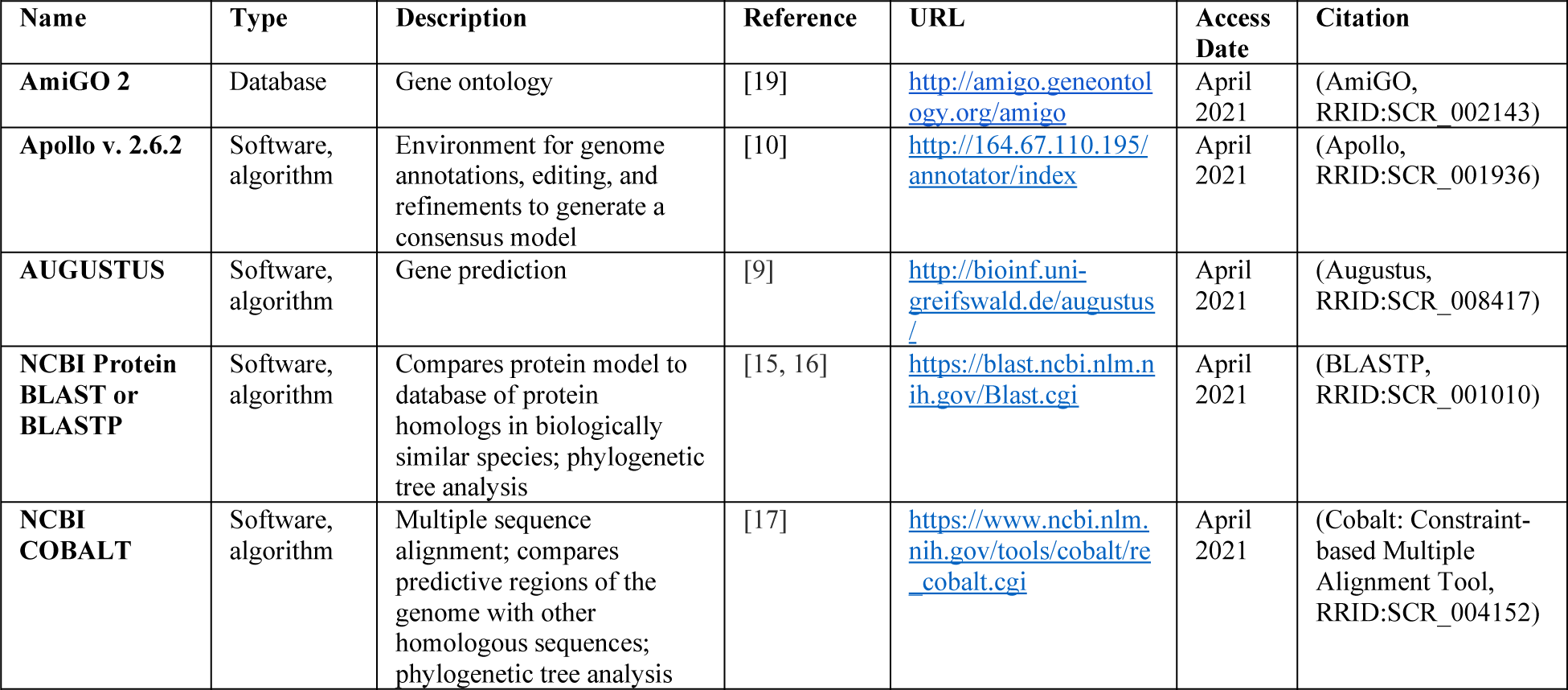
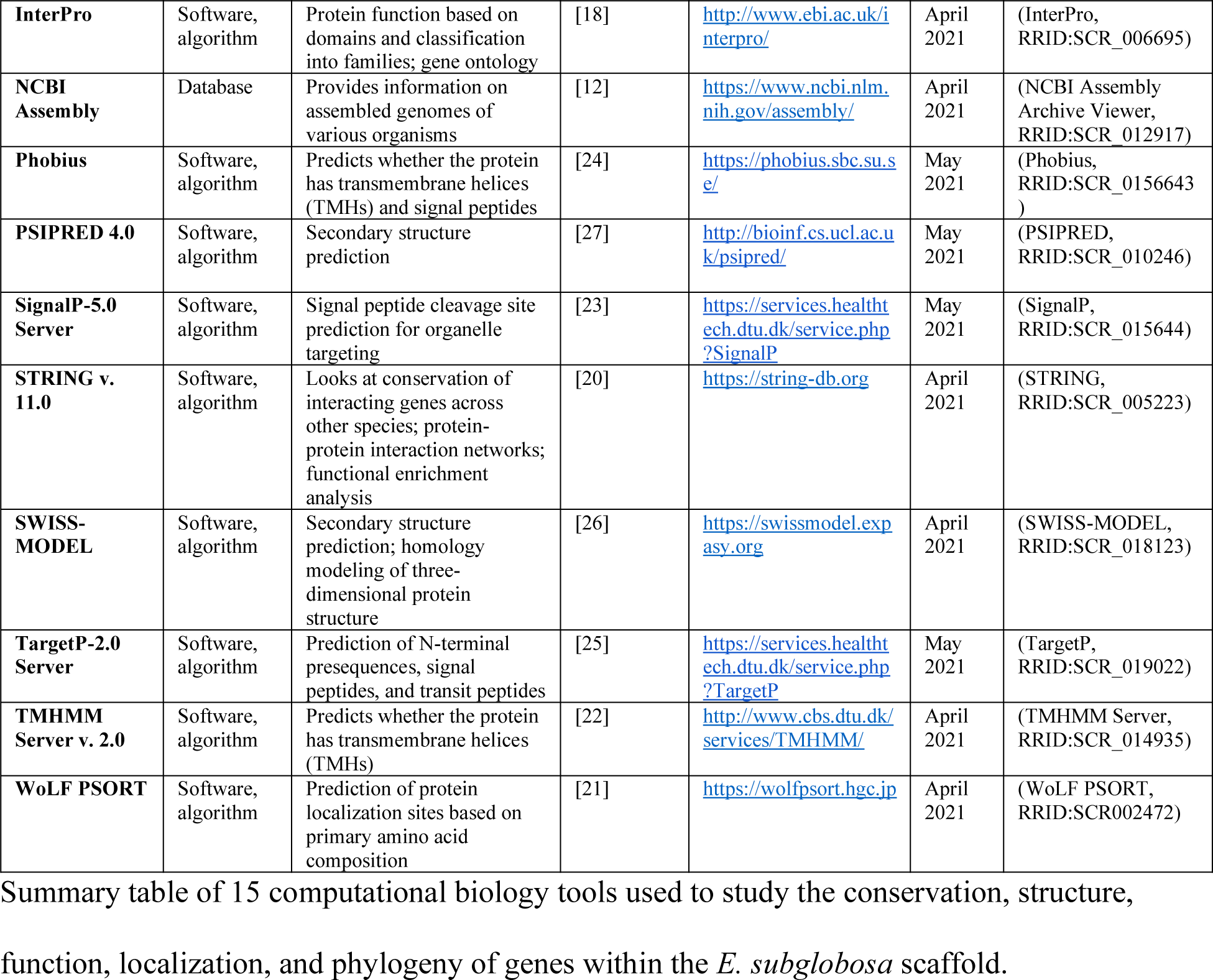
Computational biology tools.

## Results

### Mitochondrial Cholesterol Side-Chain Cleavage Enzyme

The first gene model analyzed was the g19.t1 gene located within the ML679947.1 scaffold. The AUGUSTUS gene prediction contained 520 residues and 9 exons (Fig 1A). Local pairwise alignment on BLAST showed a high query coverage of approximately 100%, high percent identity ranging from 83-87%, and low E value of 0, suggesting a very low probability of alignments to be obtained by chance (Fig 1B). We can be confident that this is a real alignment and that these species evolved from a common ancestor. The graphical distribution of top 10 BLAST hits also aligned with the aforementioned results, showing a query coverage across the entire gene and high conservation (Fig 1C). Through the COBALT multiple sequence alignment tool, the peptide sequence was aligned and had high conservation and similarity across other homologous proteins (Fig 1D). The predicted gene model was missing 3 residues from positions 488-490 when comparing the query sequence to the homologous sequences (Fig 1D). The two exons in Query_10009 and Query_10011 were not edited because those extra exons were specific to that sequence and not found in any others (Fig 1D). We could not find the sequence of the missing region nor the nearby region in the gene model, suggesting that a deletion occurred through evolution of *E. subglobosa*. Therefore, we concluded that AUGUSTUS predicted the most refined consensus model.

**Fig 1.**
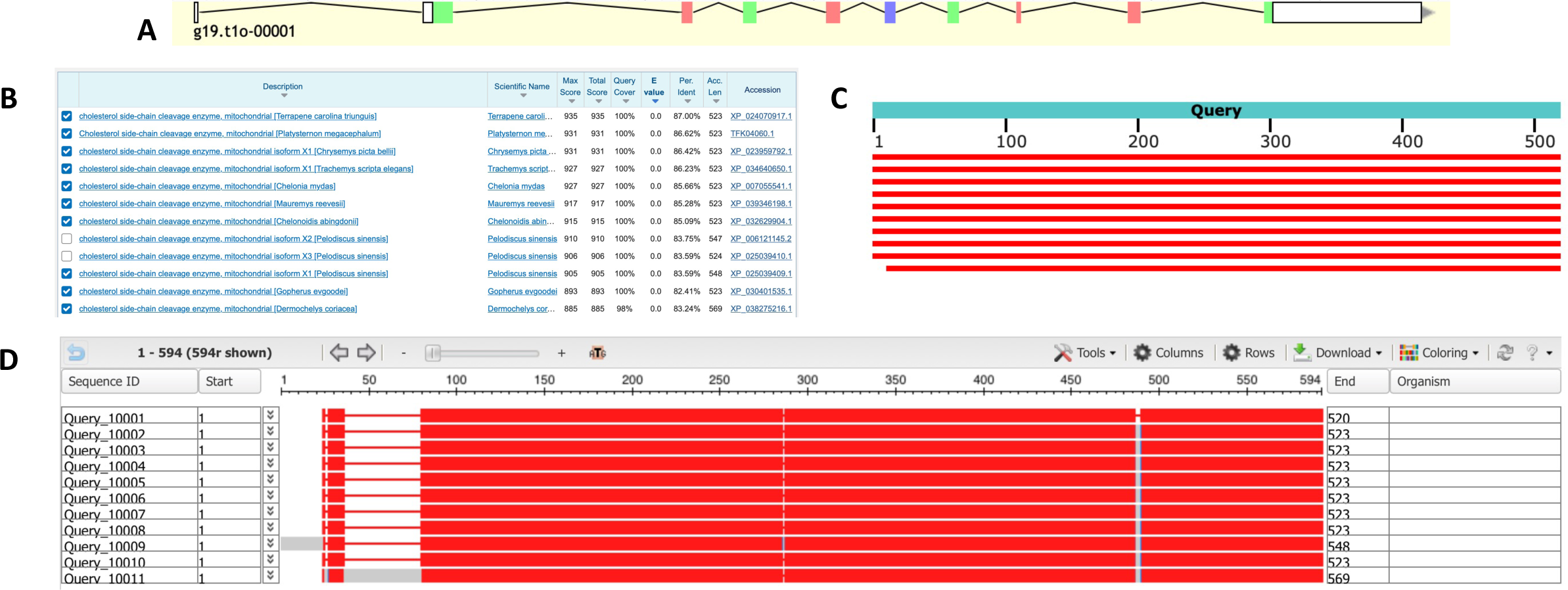
Homology-based genome annotation of the cholesterol side-chain cleavage enzyme. (A) Apollo gene editor view and AUGUSTUS track of the g19.t1 gene located within the ML679947.1 scaffold. (B) BLAST output with very high query coverage and percent identity and low E value. Top hits were all mitochondrial cholesterol side-chain cleavage enzymes. (C) Graphical representation of query coverage across the top 10 BLAST hits on 10 subject sequences. Red means high conservation. (D) COBALT multiple sequence alignment demonstrating high conservation (red) across the homologs. Low conservation is colored gray. Exons (thick lines) and introns (thin lines) are shown. Query sequence is the top, while the subjects are below.

After validation of the gene model, we analyzed the predicted function of this peptide sequence through InterPro and STRING. InterPro predicted that the protein was a mitochondrial cholesterol side-chain cleavage enzyme (Fig 2A), which concurred with BLAST (Fig 1B). The protein belonged to three families: cytochrome P450 family (IPR001128), CYP11A1 family (IPR033283), and cytochrome P450 E-class group I family (IPR002401, Fig 2A). The predicted homologous superfamily was the cytochrome P450 superfamily (IPR036396), and there was also a conserved site from residues 457-466 (IPR036396, Fig 2A). Under the GO terms, the protein was predicted to be involved in the biological process of C21-steroid hormone biosynthetic processes (GO:0006700, Fig 2B). Its predicted molecular functions included oxidoreductase activity (GO:0016705), heme binding (GO:0020037), monooxygenase activity (GO:0004497), iron ion binding (GO:0005506), and cholesterol monooxygenase (side-chain-cleaving) activity (GO:0008386, Fig 2B). Its predicted cellular component was the mitochondrion (GO:0005739, Fig 2B).

**Figure 2.**
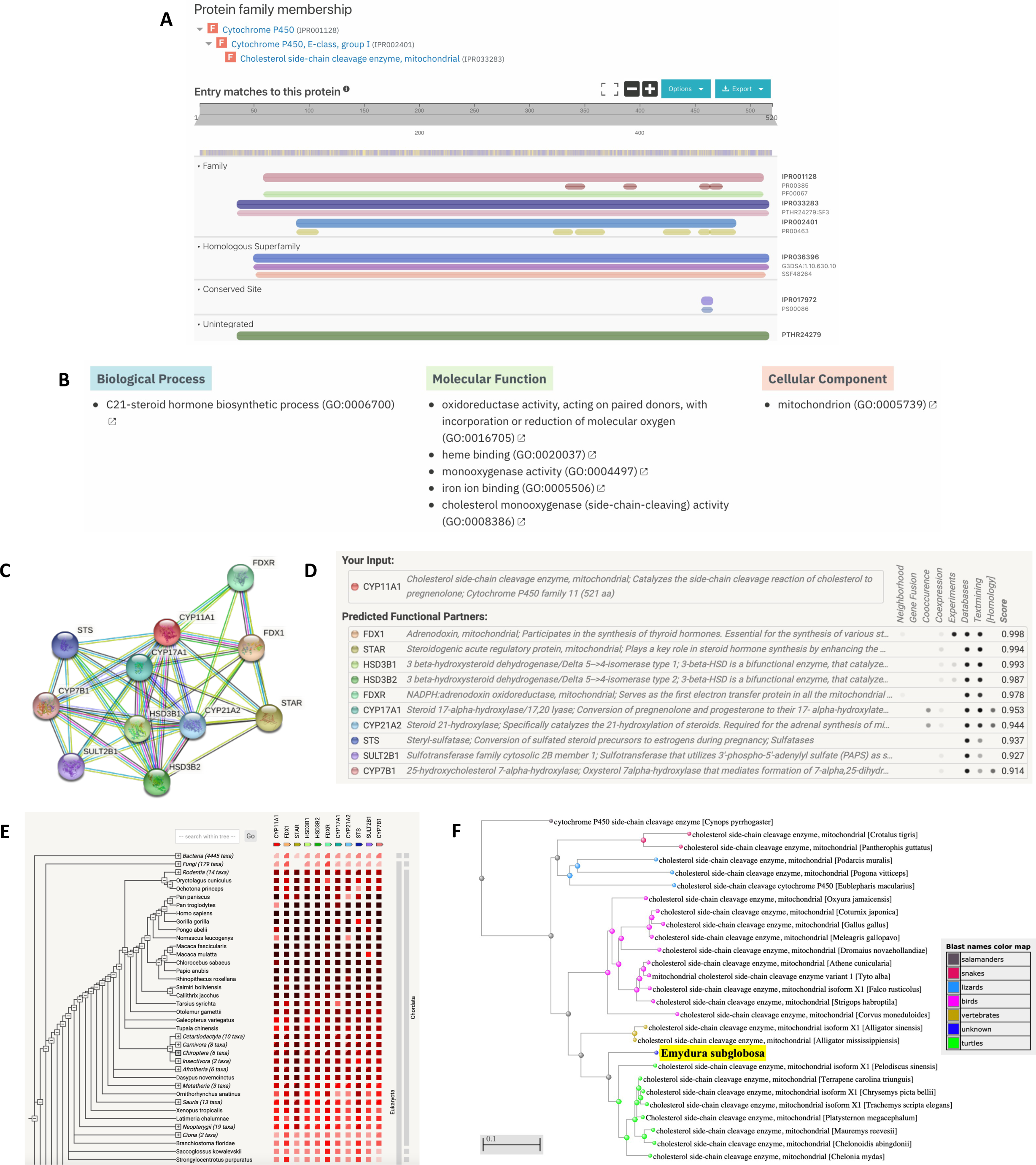
Functional analysis of the cholesterol side-chain cleavage enzyme. (A) InterPro functional analysis of the enzyme. (B) GO terms for the enzyme outputted by InterPro. (C) STRING network of predicted protein-protein interactions in *Homo sapiens*. (D) List of functional partners predicted by STRING corresponding to C. (E) Gene co-occurrence of the protein. (F) BLAST phylogenetic tree built based on pairwise alignment.

To further analyze function, we also determined protein-protein interactions through the STRING database. The protein was predicted to be CYP11A1, which plays a role in catalyzing the side-chain cleavage reaction of cholesterol in the mitochondria (Figs 2C and 2D); this agreed with the InterPro protein family results (Fig 2A). The protein’s predicted functional partners included FDX1 (a mitochondrial adrenodoxin that synthesizes thyroid hormones), STAR (a mitochondrial steroidogenic acute regulatory protein), HSD3B1/2 (dehydrogenase enzymes), and several others (Figs 2C and 2D). These network links suggest the co-evolution of the non-homologous proteins since they are functionally related and could warrant further analysis. The interaction between CYP11A1 and FDX1 was determined experimentally, while the interaction between CYP11A1 and CYP17A1, CYP21A2, and CYP7B1 were determined through homology (Fig 2D). Further research can be done to determine whether they act in a complex or directly bind each other. Additionally, the gene co-occurrence predicted that CYP11A1 interacts with CYP17A1 and CYP21A2, which suggests that these gene families have strong homology across the chordata phylum (Fig 2E). Finally, based on phylogenetic tree analysis of the mitochondrial cholesterol side-chain cleavage enzyme, *E. subglobosa* was an outgroup to other turtle species, suggesting that these species came from a common ancestor (Fig 2F and S1 Fig).

We also wanted to determine the subcellular localization of the protein based on any transmembrane helices (TMHs), signal sequences/peptides, and transit peptides. In WoLF-PSORT, a large majority of the highest matches predicted a mitochondrial localization, suggesting that this protein is likely located in the mitochondria (Fig 3A). There were no TMHs predicted, so the protein presumably does not localize on or near the plasma membrane (Fig 3B). The protein did not have any predicted signal peptides (Fig 3C). The Phobius predictions also agreed with both TMHMM and SignalP (Fig 3D). Despite having no signal peptide to target the protein to the mitochondria, there was statistically significant evidence of a mitochondrial transfer peptide at positions 36-37 (P = 0.8743, Fig 3E). Based on the localization analysis, several pieces of evidence agree that the protein localizes to the mitochondria.

**Figure 3.**
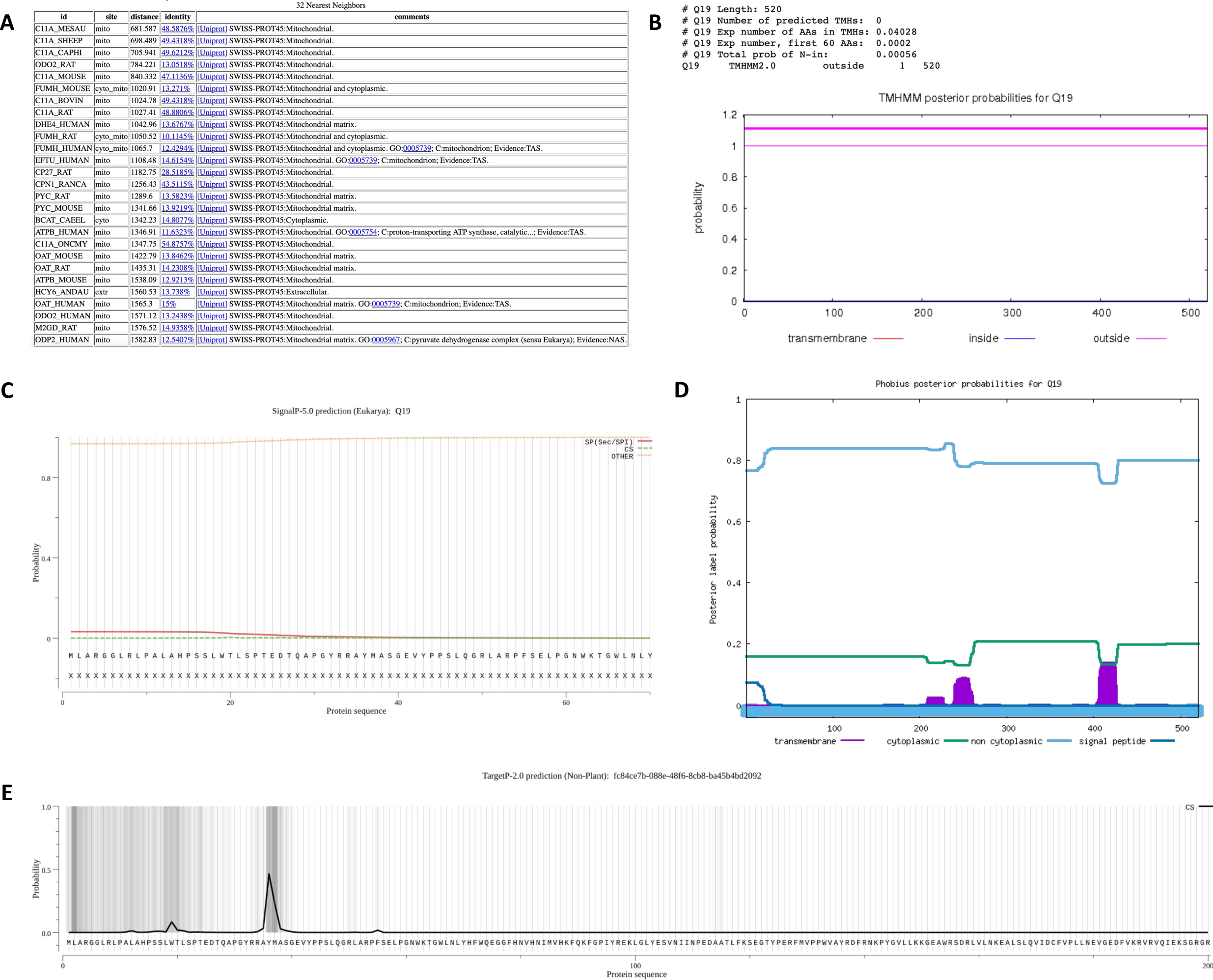
Subcellular localization of the cholesterol side-chain cleavage enzyme. (A) WoLF-PSORT prediction of the protein localization site based on 32 nearest neighbors. Localization scores are ranked from the highest scoring matches (top) to lowest (bottom). (B) TMHMM prediction of transmembrane helices (TMHs). X-axis represents the amino acid number, and y-axis represents the probability that the amino acid is located within, outside, or inside the membrane. Probabilities >0.75 are significant. (C) SignalP analysis of signal sequences existing in the amino acid sequence of the polypeptide. (D) Phobius predictions of TMHs and signal peptides. X-axis represents the amino acid number, and y-axis represents the probability that the amino acid is transmembrane, cytoplasmic, non-cytoplasmic, and/or a signal peptide. Probabilities >0.75 are significant. (E) TargetP-2.0 prediction of N-terminal pre-sequences, signal peptides, and transit peptides.

After characterizing the function and localization of the enzyme, we also wanted to understand its structure since protein structure heavily influences protein function. SWISS-MODEL predicted a monomeric structure that was highly conserved based on homology (Figs 4A and 4B). To validate the model, we examined the global and local quality estimates. The GMQE was 0.74, QMEAN was -1.77, and sequence identity was 54.04% (Fig 4B). Additionally, the local quality estimates had high probabilities in a majority of the regions; areas that dipped below 0.6 may have had poor resolution or were less conserved (Fig 4C). Comparing the model to the non-redundant set of PDB structures, the QMEAN was relatively near the average QMEAN scores for proteins of similar size (Fig 4D). The Ramachandran Plot also showed high favorability of the residues having a specific orientation, as shown by a majority of dots (95.91%) located in the dark green regions (Figs 4E and 4F). The MolProbity Score was near 0; the clash score, outliers, deviations, and bad angles were low (Fig 4F). Based on PSIPRED, there were both alpha-helixes and beta strands in the secondary structure, which was consistent with SWISS-MODEL (Fig 4G). Together, these data suggest a high-quality structure that could be used to model the mitochondrial cholesterol side-chain cleavage enzyme.

**Figure 4.**
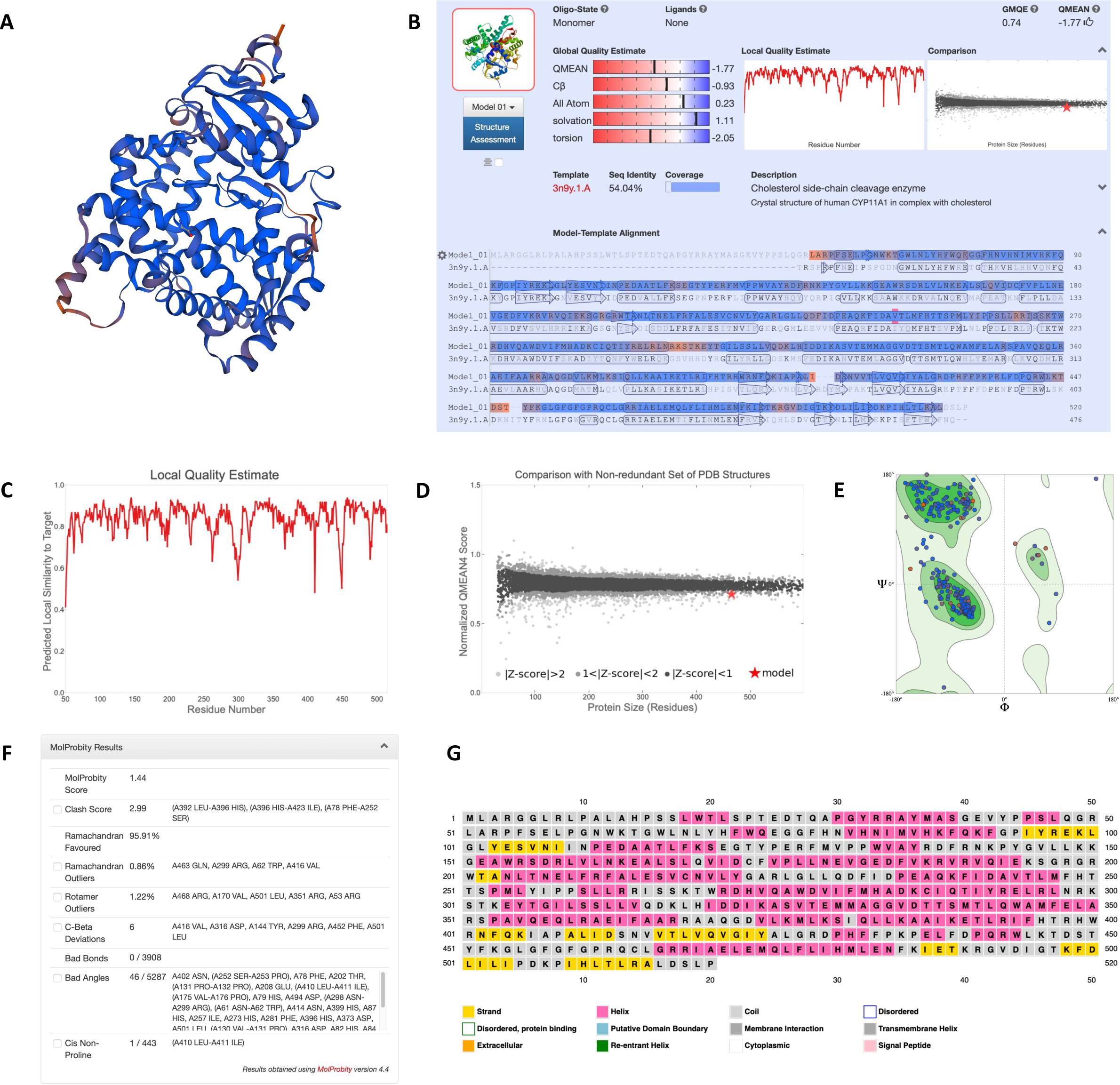
Homology modeling and structural predictions of the mitochondrial cholesterol side-chain cleavage enzyme. (A) Three-dimensional homology model built by SWISS-MODEL. Blue regions are highly conserved, while orange regions are less conserved. (B) Oligo-state, ligands, global quality estimates, template, sequence identity, and coverage outputted by SWISS-MODEL. (C) Local quality estimate showing pair residue estimates. Similarities >0.6 are high-quality models. (D) Comparison with non-redundant set of PDB structures showing QMEAN scores for experimental structures that have been deposited of similar size. The red star is our model. (E) Ramachandran plot showing the probability of a residue having a specific orientation. Dots in the dark green regions represents high probability and a high-quality model. (F) MolProbity results to validate the model. A MolProbity Score close to 0 represents the resolution that one would see a structure of this quality. Clash score represents overlapping residues; a lower value is favored. Outliers represent values that extend outside the standard deviation; low values are also favored. Low values for bad bonds and angles are also favored. (G) Secondary structure prediction through PSIPRED.

### Receptor for Retinol Uptake STRA6

The second gene model analyzed was the g24.t1 gene located within the ML679947.1 scaffold. The initial gene prediction contained 683 residues and 18 exons (Fig 5A). Local pairwise alignment on BLAST showed a high query coverage of approximately 100%, high percent identity ranging from 80-85%, and low E value of 0, suggesting a very low probability of alignments to be obtained by chance (Fig 5B). We can be confident that this alignment did not occur by chance and that these species evolved from a common ancestor. The graphical distribution of top 9 BLAST hits also aligns with the aforementioned results, showing a query coverage across the entire gene and high conservation (Fig 5C). Through the COBALT multiple sequence alignment tool, the peptide sequence was aligned and had high conservation and similarity across other homologous proteins (Fig 5D). The predicted gene model had 17 extra residues from positions 263-279 when comparing the query sequence to the homologous sequences (Fig 5D). The exon in Query_10008 was not edited because it was specific to that sequence and not found in any others (Fig 5D). The 17 residues spanned across an entire exon (Fig 5E), so the entire exon was removed from the gene prediction. Therefore, the most refined consensus model contained 666 residues and 17 exons (Fig 5A). BLAST results after gene editing revealed a higher percent identity ranging from 82-87% (Fig 5B), and COBALT alignment was highly conserved across the gene (Fig 5D).

**Figure 5.**
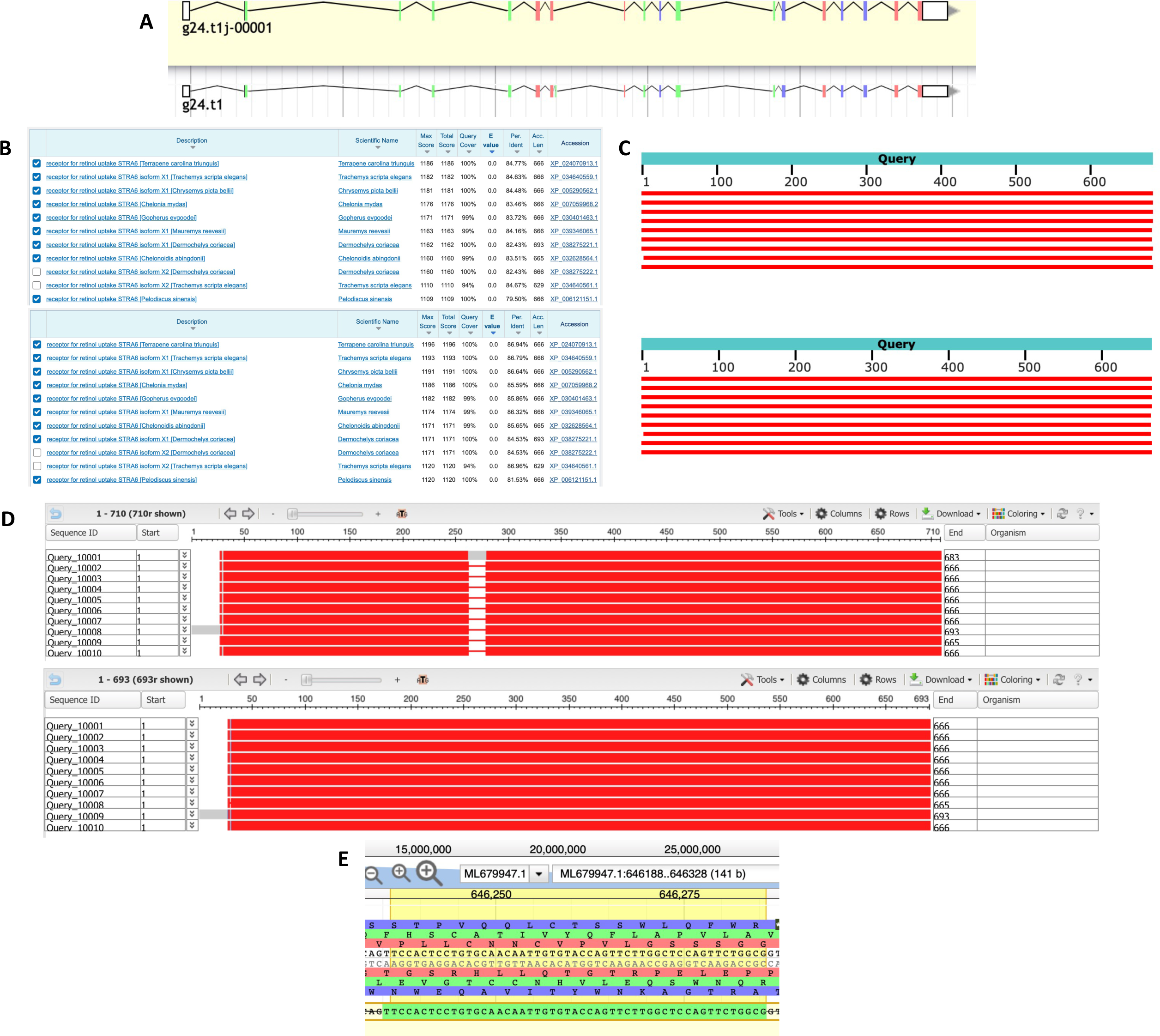
Homology-based genome annotation of the receptor for retinol uptake STRA6. (A) Apollo gene editor view and AUGUSTUS track of the g24.t1 gene located within the ML679947.1 scaffold. Bottom: initial *ab initio* prediction. Top: consensus gene model. (B) BLAST output before (top) and after (bottom) genome editing with very high query coverage and percent identity and low E value. Top hits were all receptors for retinol uptake STRA6. (C) Graphical representation of query coverage across the top 9 BLAST hits on 9 subject sequences before (top) and after (bottom) genome editing. Red means high conservation. (D) COBALT multiple sequence alignment before (top) and after (bottom) genome editing, demonstrating high conservation (red) across the homologs. Low conservation is colored gray. Exons (thick lines) and introns (thin lines) are shown. Query sequence is the top, while the subjects are below. (E) Exon containing 17 extra residues at positions 263-279.

After validation of the gene model, we analyzed the predicted function of this peptide sequence through InterPro and STRING. InterPro predicted that the protein was a receptor for retinol uptake STRA6 (Fig 6A), which concurred with BLAST (Fig 5B). The protein belonged to the STRA6 family (IPR026612) and contained various predictions of non-cytoplasmic, cytoplasmic, and transmembrane domains (Fig 6A). Under the GO terms, its predicted molecular functions included retinol transmembrane transporter activity (GO:0034632) and signaling receptor activity (GO:0038023, Fig 6B).

**Figure 6.**
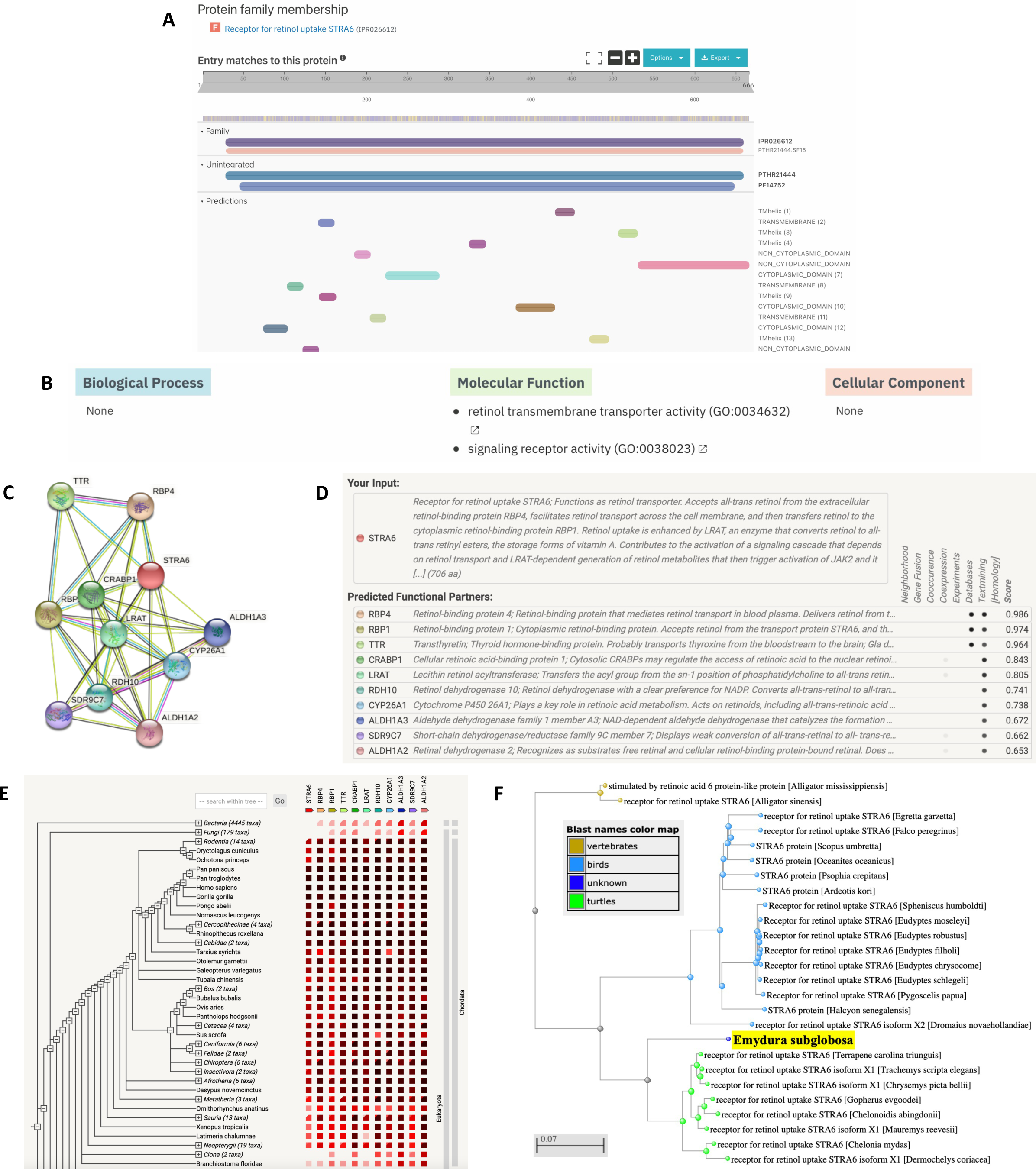
Functional analysis of the receptor for retinol uptake STRA6. (A) InterPro functional analysis of the receptor protein. (B) GO terms for the protein outputted by InterPro. (C) STRING network of predicted protein-protein interactions in *Homo sapiens*. (D) List of functional partners predicted by STRING corresponding to C. (E) Gene co-occurrence of the protein. (F) BLAST phylogenetic tree built based on pairwise alignment.

To further analyze function, we also determined protein-protein interactions through the STRING database. The protein was predicted to be STRA6, which plays a role in facilitating retinol transport across the cell membrane and contributes to a retinol-dependent signaling cascade (Figs 6C and 6D); this agreed with the InterPro protein family results (Fig 6A). The protein’s predicted functional partners included RBP1/4 (retinol binding proteins that mediate retinol transport), TTR (a thyroid hormone-binding protein), CYP26A1 (cytochrome P450 that aids retinoic acid metabolism), and several others (Figs 6C and 6D). These network links suggest the co-evolution of the non-homologous proteins since they are functionally related and could warrant additional analysis. Further research can also be done to determine whether they act in a complex or directly bind each other. Additionally, the gene co-occurrence suggests that these gene families have strong homology across the chordata phylum (Fig 6E). Finally, based on phylogenetic tree analysis of the STRA6 receptor, *E. subglobosa* was an outgroup to other turtle species, suggesting that these species came from a common ancestor (Fig 6F and S2 Fig).

We also wanted to analyze the subcellular localization of the protein based on any TMHs, signal sequences/peptides, and transit peptides. In WoLF-PSORT, all of the highest matches predicted an integral membrane protein, suggesting that this protein is likely located in the plasma membrane (Fig 7A). There were 10 TMHs predicted (residues 52-74, 104-123, 143-162, 199-221, 291-313, 326-345, 360-382, 431-453, 473-495, and 508-530), so the protein shows a high likelihood of localizing to the plasma membrane (Fig 7B). The protein was predicted to not have any signal peptides or transfer peptides (Figs 7C and 7E). The Phobius predictions also agreed with both TMHMM and SignalP (Fig 7D). Based on the localization analysis, there is strong evidence supporting that STRA6 localizes to the plasma membrane as an integral membrane protein.

**Figure 7.**
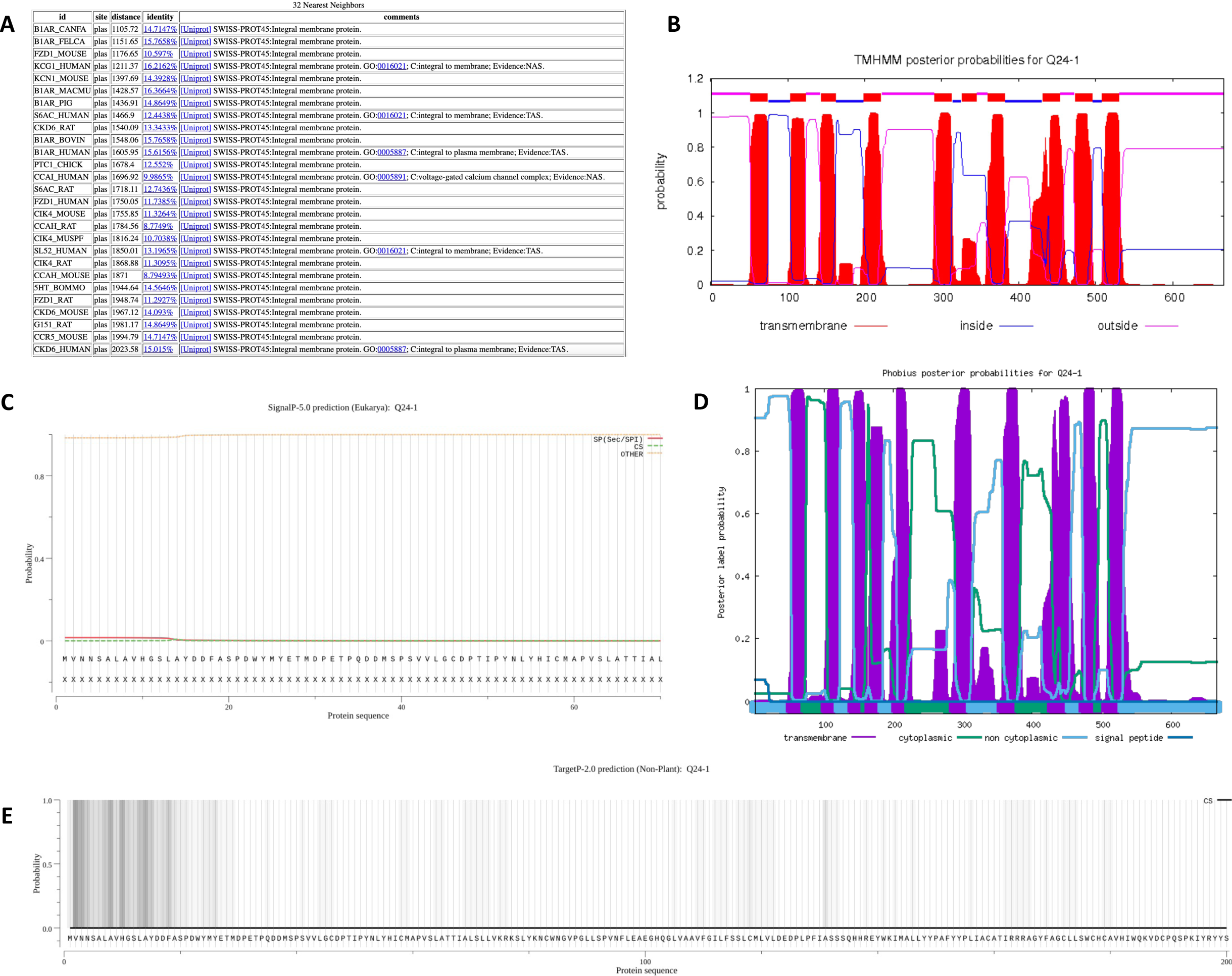
Subcellular localization of the receptor for retinol uptake STRA6. (A) WoLF-PSORT prediction of the protein localization site based on 32 nearest neighbors. Localization scores are ranked from the highest scoring matches (top) to lowest (bottom). (B) TMHMM prediction of TMHs. X-axis represents the amino acid number, and y-axis represents the probability that the amino acid is located within, outside, or inside the membrane. Probabilities >0.75 are significant. (C) SignalP analysis of signal sequences existing in the amino acid sequence of the polypeptide. (D) Phobius predictions of TMHs and signal peptides. X-axis represents the amino acid number, and y-axis represents the probability that the amino acid is transmembrane, cytoplasmic, non-cytoplasmic, and/or a signal peptide. Probabilities >0.75 are significant. (E) TargetP-2.0 prediction of N-terminal pre-sequences, signal peptides, and transit peptides.

After identifying the function and localization of the enzyme, we also wanted to understand its structure since protein structure heavily influences protein function. The SWISS-MODEL predicted a monomeric structure that was somewhat conserved based on homology (Figs 8A and 8B). To validate the model, we examined the global and local quality estimates. The GMQE was 0.67, QMEAN was -5.64, and sequence identity was 47.65% (Fig 8B). Additionally, the local quality estimates had a mix of high and low probabilities in various positions; areas that dipped below 0.6 may have had poor resolution or were less conserved (Fig 8C). Comparing the model to the non-redundant set of PDB structures, the QMEAN was much lower than the average QMEAN scores for proteins of similar size (Fig 8D). However, the Ramachandran Plot also showed high favorability of the residues having a specific orientation, as shown by a majority of dots (91.30%) located in the dark green regions (Figs 8E and 8F). The MolProbity Score was near 0; the clash score, outliers, deviations, and bad angles were low (Fig 8F). PSIPRED predicted only alpha-helixes in the secondary structure, which was consistent with SWISS-MODEL (Fig 8G). Based on these contradicting data, we do not have confidence in the model, as it diverges from the actual protein structure. If we determined the 3D structure experimentally, it would be likely be different from SWISS-MODEL.

**Figure 8.**
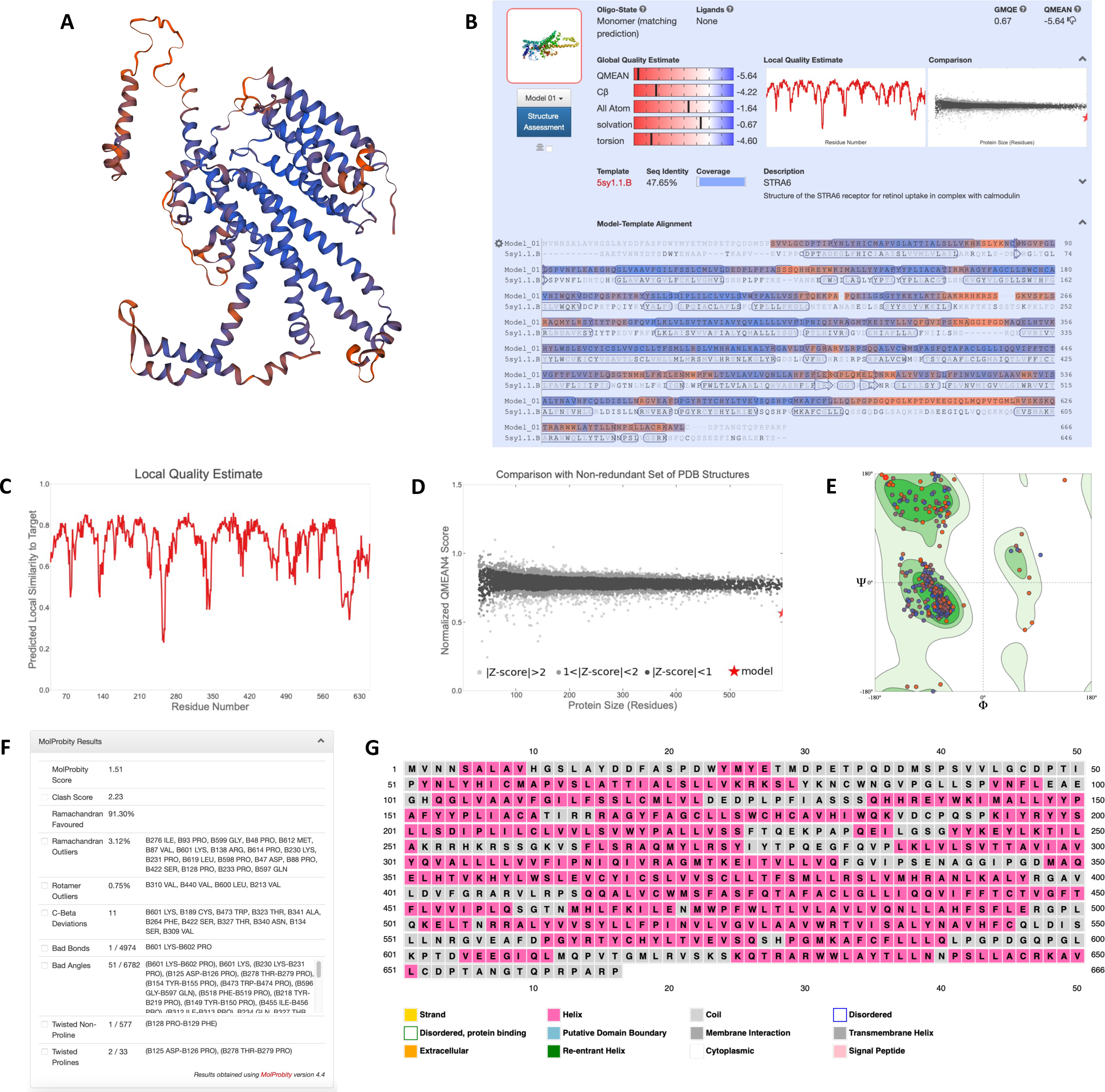
Homology modeling and structural predictions of the receptor for retinol uptake STRA6. (A) Three-dimensional homology model built by SWISS-MODEL. Blue regions are highly conserved, while orange regions are less conserved. (B) Oligo-state, ligands, global quality estimates, template, sequence identity, and coverage outputted by SWISS-MODEL. (C) Local quality estimate showing pair residue estimates. Similarities >0.6 are high-quality models. (C) Comparison with non-redundant set of PDB structures showing QMEAN scores for experimental structures that have been deposited of similar size. The red star is our model. (E) Ramachandran plot showing the probability of a residue having a specific orientation. Dots in the dark green regions represents high probability and a high-quality model. (F) MolProbity results to validate the model. A MolProbity Score close to 0 represents the resolution that one would see a structure of this quality. Clash score represents overlapping residues; a lower value is favored. Outliers represent values that extend outside the standard deviation; low values are also favored. Low values for bad bonds and angles are also favored. (G) Secondary structure prediction through PSIPRED.

### Lysyl Oxidase Homolog 1

The third gene model analyzed was the g32.t1 gene located within the ML679947.1 scaffold. The AUGUSTUS gene prediction contained 638 residues and 8 exons (Fig 9A). Local pairwise alignment on BLAST showed a high query coverage of 95%, high percent identity ranging from 81-87%, and low E value of 0, suggesting a very low probability of alignments to be obtained by chance (Fig 9B). We can be confident that this is a real alignment and that these species evolved from a common ancestor. The graphical distribution of top 10 BLAST hits also aligns with the aforementioned results, showing a query coverage across the entire gene and high conservation (Fig 9C). Through the COBALT multiple sequence alignment tool, the peptide sequence was aligned and had high conservation and similarity across other homologous proteins (Fig 9D). When comparing the query sequence to the homologous sequences, the predicted gene model was missing 3 residues from positions 15-17 and 3 residues from positions 347-349 (Fig 9D). It also had 2 extra residues from positions 258-259, 4 extra residues from positions 405-408, and 30 extra residues from positions 630-659. For the two regions with missing sequences, we could not find them in the middle of the exon (Fig 9E), suggesting that there were deletions that occurred through evolution of *E. subglobosa*. We removed the extra residues by splitting or deleting the exon. Therefore, the most refined consensus model contained 602 residues and 9 exons (Fig 9A). BLAST results after gene editing revealed a higher query coverage of 99-100% and higher percent identity ranging from 81-88% (Fig 9B), and COBALT alignment was highly conserved across the gene (Fig 9D).

**Figure 9.**
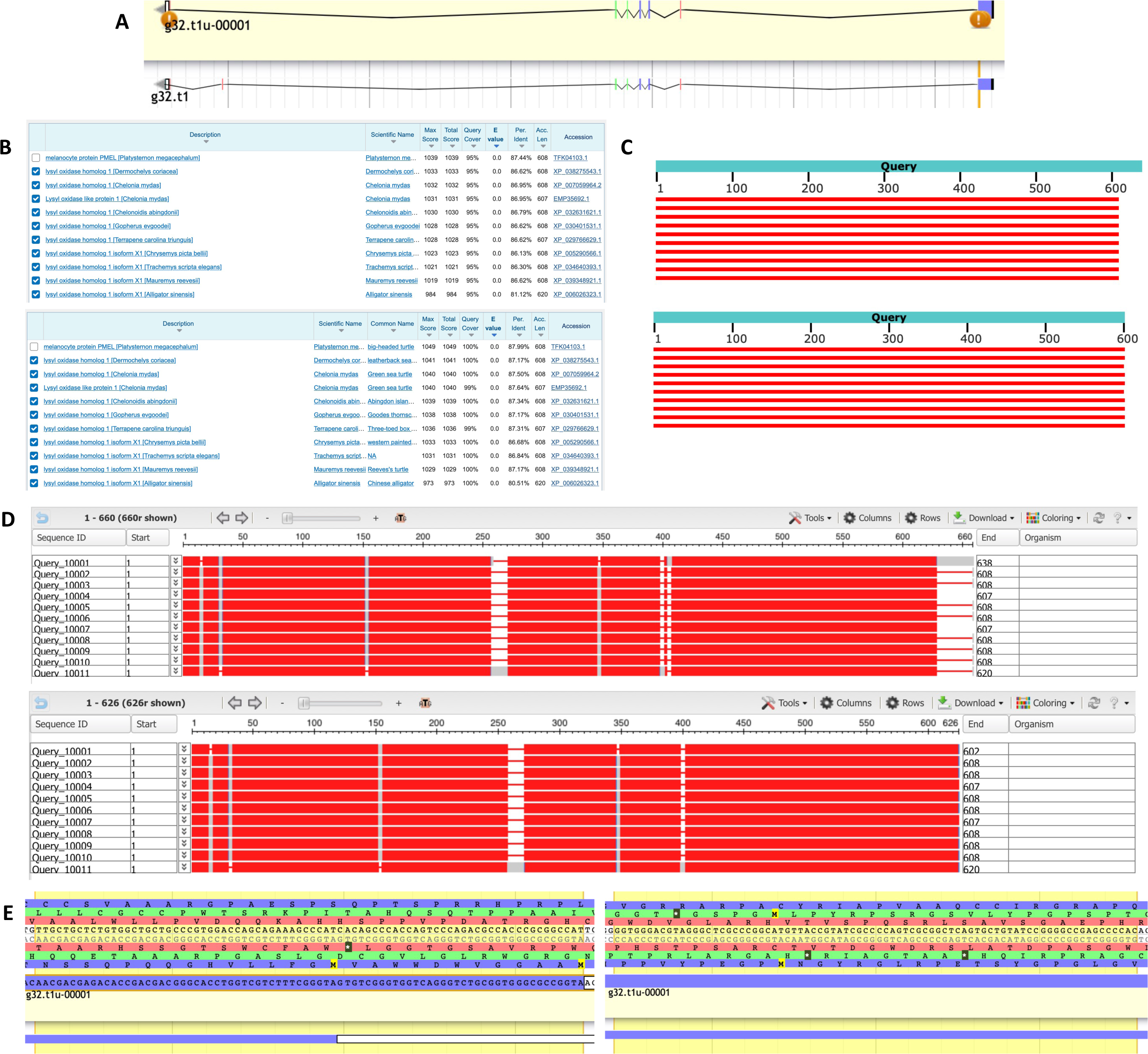
Homology-based genome annotation of lysyl oxidase homolog 1. (A) Apollo gene editor view and AUGUSTUS track of the g32.t1 gene located within the ML679947.1 scaffold. Bottom: initial *ab initio* prediction. Top: consensus gene model. (B) BLAST output before (top) and after (bottom) genome editing with very high query coverage and percent identity and low E value. Top hits were all lysyl oxidase homolog 1. (C) Graphical representation of query coverage across the top 10 BLAST hits on 10 subject sequences before (top) and after (bottom) genome editing. Red means high conservation. (D) COBALT multiple sequence alignment before (top) and after (bottom) genome editing, demonstrating high conservation (red) across the homologs. Low conservation is colored gray. Exons (thick lines) and introns (thin lines) are shown. Query sequence is the top, while the subjects are below. (E) Missing 3 residues from positions 15-17 (left) and 3 residues from positions 347-349 (right) in the middle of the exons.

After validation of the gene model, we analyzed the predicted function of this peptide sequence through InterPro and STRING. InterPro predicted that the protein was a lysyl oxidase homolog 1 (Fig 10A), which concurred with BLAST (Fig 9B). The protein belonged to the lysyl oxidase family (IPR001695) and there was also a conserved site from residues 471-484 (IPR019828, Fig 10A). Under the GO terms, its predicted molecular functions included oxidoreductase activity, acting on the CH-NH2 group of donors, oxygen as acceptor (GO:0016641) and copper ion binding (GO:0005507, Fig 10B).

**Figure 10.**
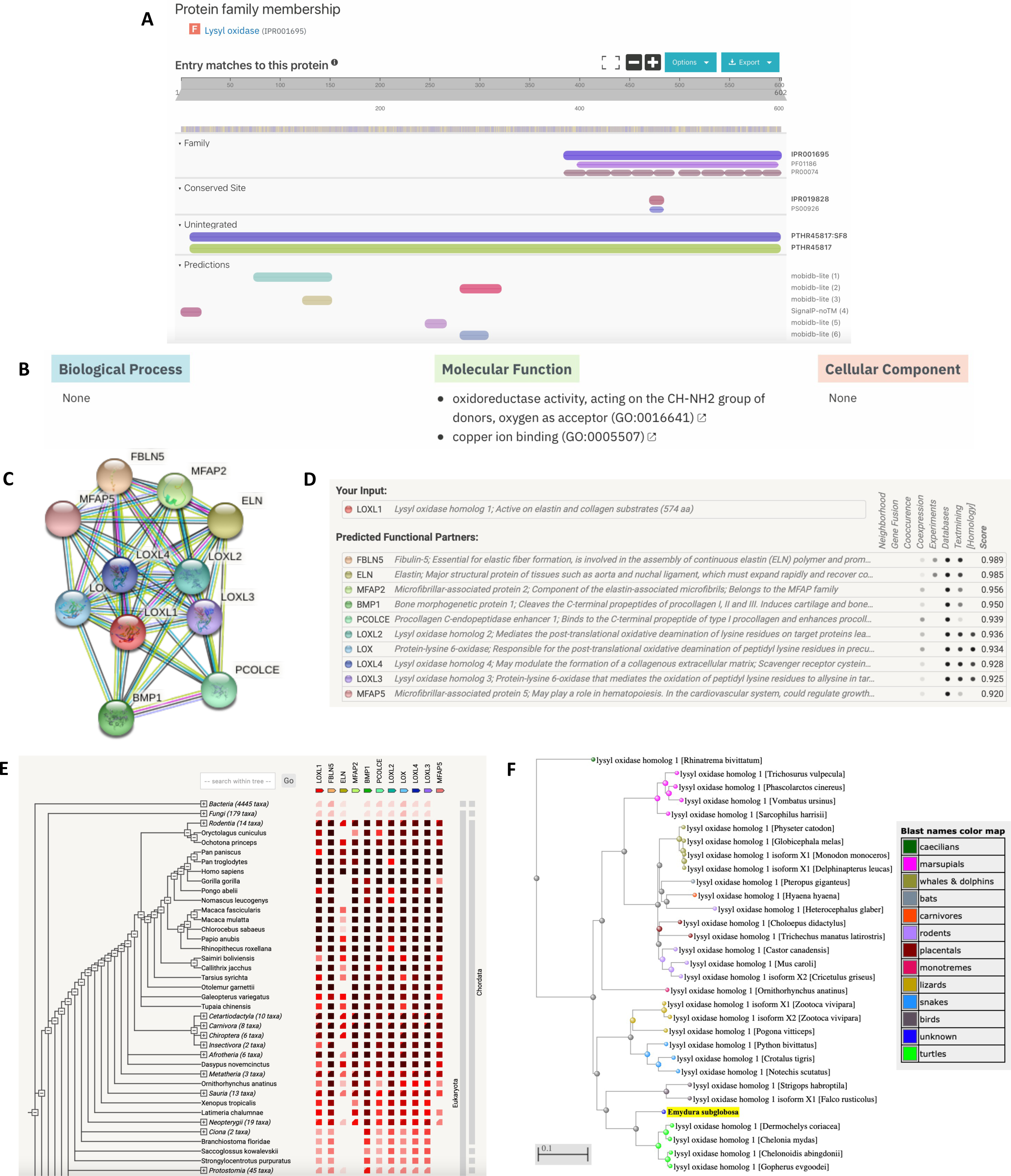
Functional analysis of lysyl oxidase homolog 1. (A) InterPro functional analysis of the enzyme. (B) GO terms for the protein outputted by InterPro. (C) STRING network of predicted protein-protein interactions in *Homo sapiens*. (D) List of functional partners predicted by STRING corresponding to C. (E) Gene co-occurrence of the enzyme. (F) BLAST phylogenetic tree built based on pairwise alignment.

To further analyze function, we also determined protein-protein interactions through the STRING database. The protein was predicted to be LOXL1, which plays a role in elastin and collagen substrates (Figs 10C and 10D); this agreed with the InterPro protein family results (Fig 10A). The protein’s predicted functional partners included FBLN5 (fibulin-5 protein involved in elastin fiber formation), ELN (elastin), BMP1 (bone morphogenetic protein that cleaves C-terminal pro-peptides of procollagen), and several others (Figs 10C and 10D). These network links suggest the co-evolution of the non-homologous proteins since they are functionally related and could warrant additional analysis. Further research can also be done to determine whether they act in a complex or directly bind each other. Additionally, the gene co-occurrence suggests that these gene families have strong homology across the chordata phylum (Fig 10E). Finally, based on phylogenetic tree analysis of the lysyl oxidase homolog 1, *E. subglobosa* was an outgroup to other turtle species, suggesting that these species came from a common ancestor (Fig 10F and S3 Fig).

We also wanted to determine the subcellular localization of the protein based on any TMHs, signal sequences/peptides, and transit peptides. In WoLF-PSORT, there was a variety of locations predicted by the highest matches, but the most abundant were the nucleus and extracellular regions (Fig 11A). There were no TMHs predicted (Fig 11B) and only non-cytoplasmic domains (Fig 11D), so the protein likely does not localize to the plasma membrane. The protein was predicted to have a signal peptide cleavage site between residues 20-21 (P = 0.8926, Figs 10C and 10E). Based on the data, lysyl oxidase homolog 1 is predicted to localize to the nucleus and/or extracellular regions and does not localize to the plasma membrane.

**Figure 11.**
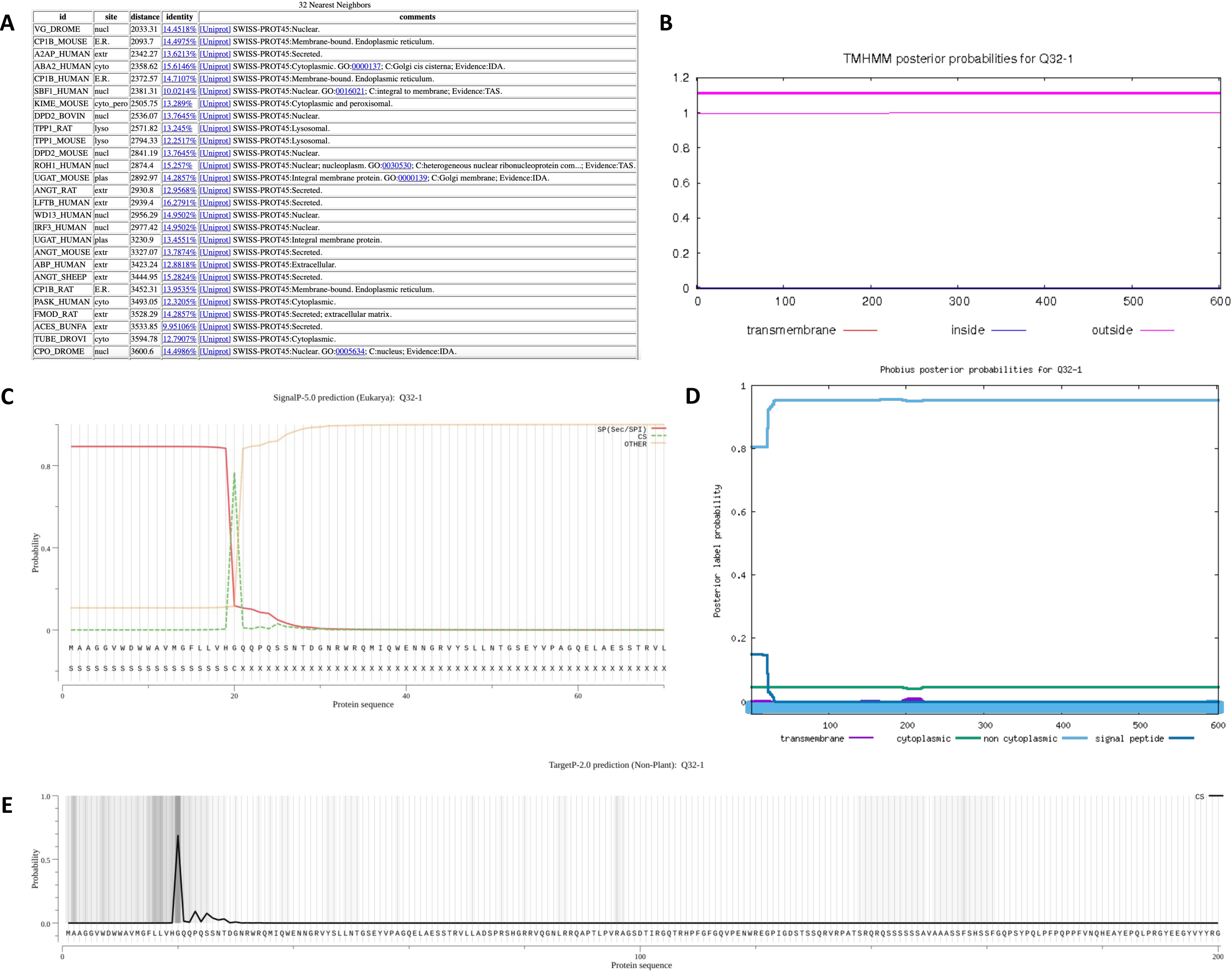
Subcellular localization of lysyl oxidase homolog 1. (A) WoLF-PSORT prediction of the protein localization site based on 32 nearest neighbors. Localization scores are ranked from the highest scoring matches (top) to lowest (bottom). (B) TMHMM prediction of TMHs. X-axis represents the amino acid number, and y-axis represents the probability that the amino acid is located within, outside, or inside the membrane. Probabilities >0.75 are significant. (C) SignalP analysis of signal sequences existing in the amino acid sequence of the polypeptide. (D) Phobius predictions of TMHs and signal peptides. X-axis represents the amino acid number, and y-axis represents the probability that the amino acid is transmembrane, cytoplasmic, non-cytoplasmic, and/or a signal peptide. Probabilities >0.75 are significant. (E) TargetP-2.0 prediction of N-terminal pre-sequences, signal peptides, and transit peptides.

After characterizing the function and localization of the enzyme, we also wanted to understand its structure since protein structure heavily influences protein function. The SWISS-MODEL predicted a monomeric structure that was somewhat conserved based on homology (Figs 12A and 12B). To validate the model, we examined the global and local quality estimates. The GMQE was 0.22, QMEAN was -2.59, and sequence identity was 48.53% (Fig 12B). Additionally, the local quality estimates had mostly regions of high probabilities; areas that dipped below 0.6 may have had poor resolution or were less conserved (Fig 12C). Comparing the model to the non-redundant set of PDB structures, the QMEAN was slightly below the average QMEAN scores for proteins of similar size (Fig 12D). However, the Ramachandran Plot also showed high favorability of the residues having a specific orientation, as shown by a majority of dots (90.20%) located in the dark green regions (Figs 12E and 12F). The MolProbity Score was near 0; the clash score, outliers, deviations, and bad angles were low (Fig 12F). PSIPRED prediced both alpha-helixes and beta-strands in the secondary structure, which was consistent with SWISS-MODEL (Fig 12G). Together, these data suggest a high-quality structure that could be used to model the lysyl oxidase homolog 1 enzyme.

**Figure 12.**
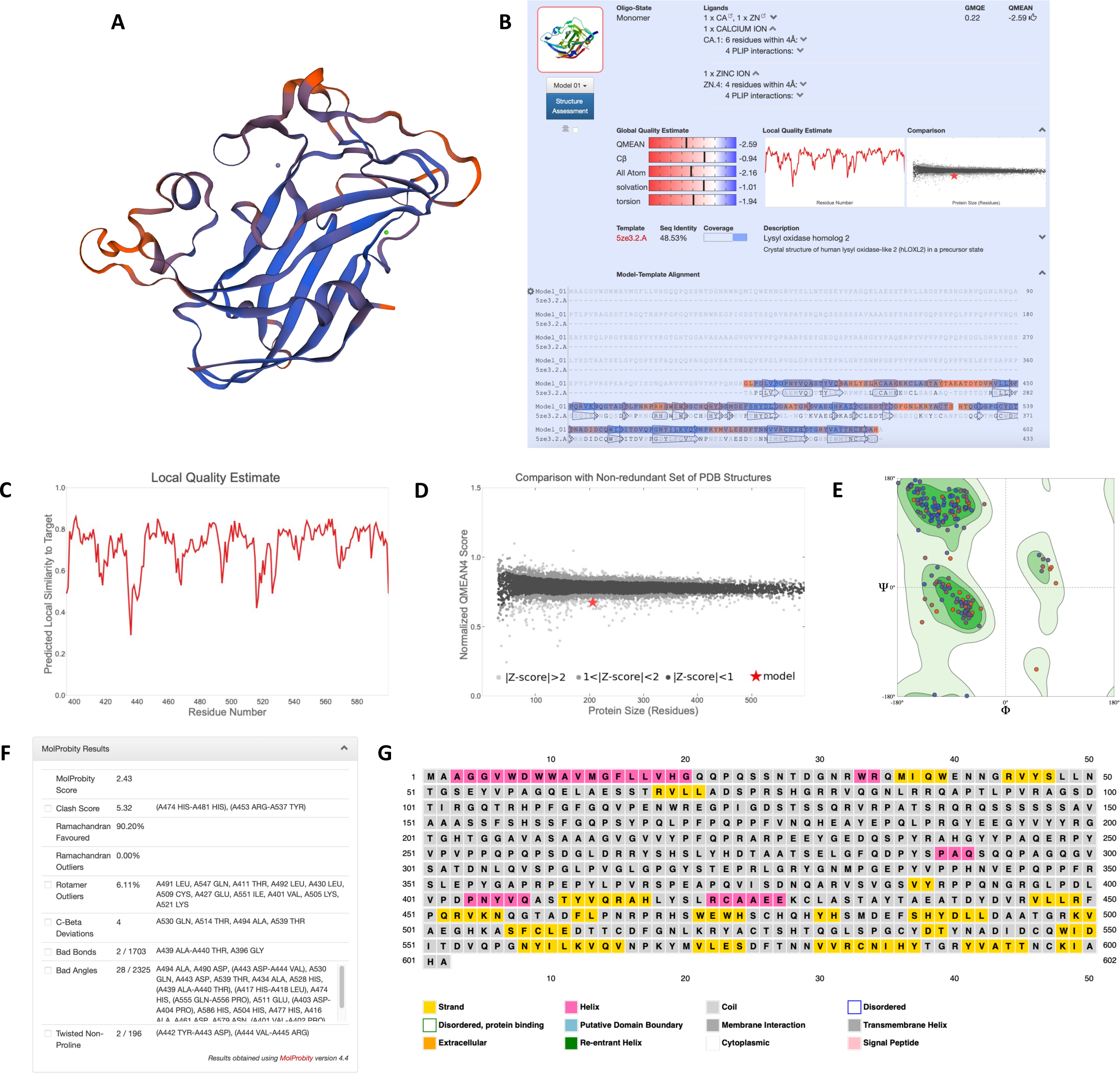
Homology modeling and structural predictions of lysyl oxidase homolog 1. (A) Three-dimensional homology model built by SWISS-MODEL. Blue regions are highly conserved, while orange regions are less conserved. (B) Oligo-state, ligands, global quality estimates, template, sequence identity, and coverage outputted by SWISS-MODEL. (C) Local quality estimate showing pair residue estimates. Similarities >0.6 are high-quality models. (D) Comparison with non-redundant set of PDB structures showing QMEAN scores for experimental structures that have been deposited of similar size. The red star is our model. (E) Ramachandran plot showing the probability of a residue having a specific orientation. Dots in the dark green regions represents high probability and a high-quality model. (F) MolProbity results to validate the model. A MolProbity Score close to 0 represents the resolution that one would see a structure of this quality. Clash score represents overlapping residues; a lower value is favored. Outliers represent values that extend outside the standard deviation; low values are also favored. Low values for bad bonds and angles are also favored. (G) Secondary structure prediction through PSIPRED.

### Mitochondrial Methylmalonyl-CoA Epimerase

The final gene model analyzed was the g112.t1 gene located within the ML679947.1 scaffold. The AUGUSTUS gene prediction contained 122 residues and 3 exons (Fig 13A). Local pairwise alignment on BLAST showed a high query coverage of 89-98%, high percent identity ranging from 55-92%, and low E values, suggesting a very low probability of alignments to be obtained by chance (Fig 13B). We can be confident that this alignment did not occur by chance and that these species evolved from a common ancestor. The graphical distribution of top 7 BLAST hits also aligns with the aforementioned results, showing a query coverage across the entire gene but relatively moderate conservation (Fig 13C). Through the COBALT multiple sequence alignment tool, the peptide sequence was aligned and had high conservation and similarity across other homologous proteins in certain areas (Fig 13D). When comparing the query sequence to the homologous sequences, the predicted gene model was missing 54 residues from positions 82-135 and had 1 extra residue at position 20 (Fig 13D). For the missing sequences, we extended the exon to encompass the missing residues in the coding region of the gene; for the extra residue, we shortened the exon by one amino acid since it was on the end of the exon. Therefore, the most refined consensus model contained 174 residues and 4 exons (Fig 13A). BLAST results after gene editing revealed a higher query coverage of 100% and higher percent identity ranging from 90-94% (Fig 13B), and COBALT alignment was highly conserved across the gene (Fig 13D).

**Figure 13.**
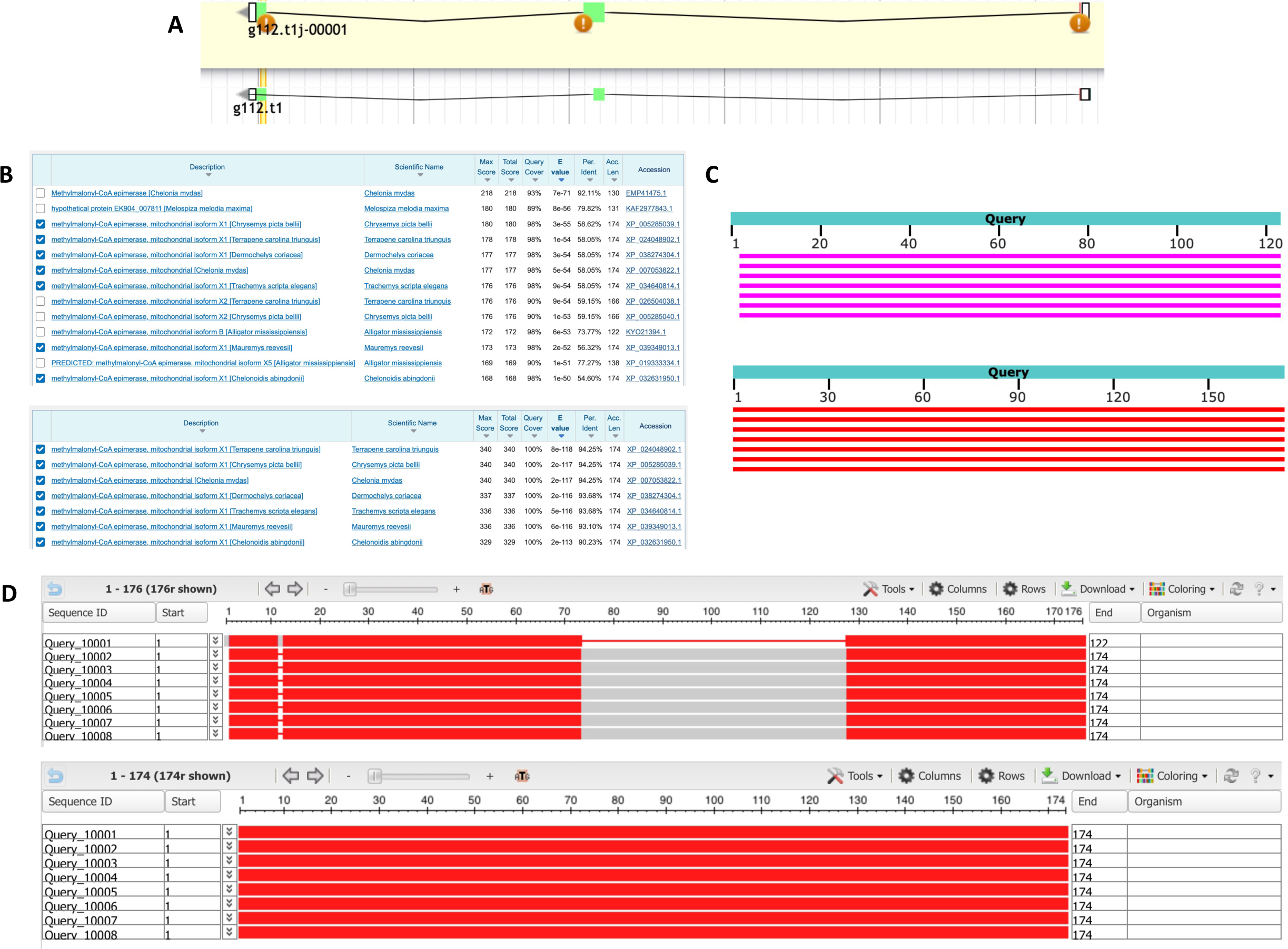
Homology-based genome annotation of the methylmalonyl-CoA epimerase (MCEE) enzyme. (A) Apollo gene editor view and AUGUSTUS track of the g112.t1 gene located within the ML679947.1 scaffold. Bottom: initial *ab initio* prediction. Top: consensus gene model. (B) BLAST output before (top) and after (bottom) genome editing with very high query coverage and percent identity and low E value. Top hits were all methylmalonyl-CoA epimerase. (C) Graphical representation of query coverage across the top 7 BLAST hits on 7 subject sequences before (top) and after (bottom) genome editing. Red means high conservation, and magenta means some conservation. (D) COBALT multiple sequence alignment before (top) and after (bottom) genome editing, demonstrating high conservation (red) across the homologs. Low conservation is colored gray. Exons (thick lines) and introns (thin lines) are shown. Query sequence is the top, while the subjects are below.

After validation of the gene model, we analyzed the predicted function of this peptide sequence through InterPro, AmiGO 2, and STRING. InterPro predicted that the protein was a methylmalonyl-CoA epimerase (Fig 14A), which concurred with BLAST (Fig 13B). The protein belonged to the methylmalonyl-CoA epimerase family (IPR017515) with metal binding and substrate binding sites and dimer interfaces (Fig 14A). The protein was also predicted to be a part of the VOC domain (IPR037523) and glyoxalase/bleomycin resistance protein/dihydroxybiphenyl dioxygenase homologous superfamily (IPR029068, Fig 14A). There were also predictions of a signal peptide and non-cytoplasmic domains (Fig 14A). No GO terms were outputted on InterPro; however, there were two GO terms outputted by AmiGO 2 which were the biological process L-methylmalonyl-CoA metabolic (GO:0046491) and the molecular function methylmalonyl-CoA epimerase (GO:0004493, S4 Fig).

**Figure 14.**
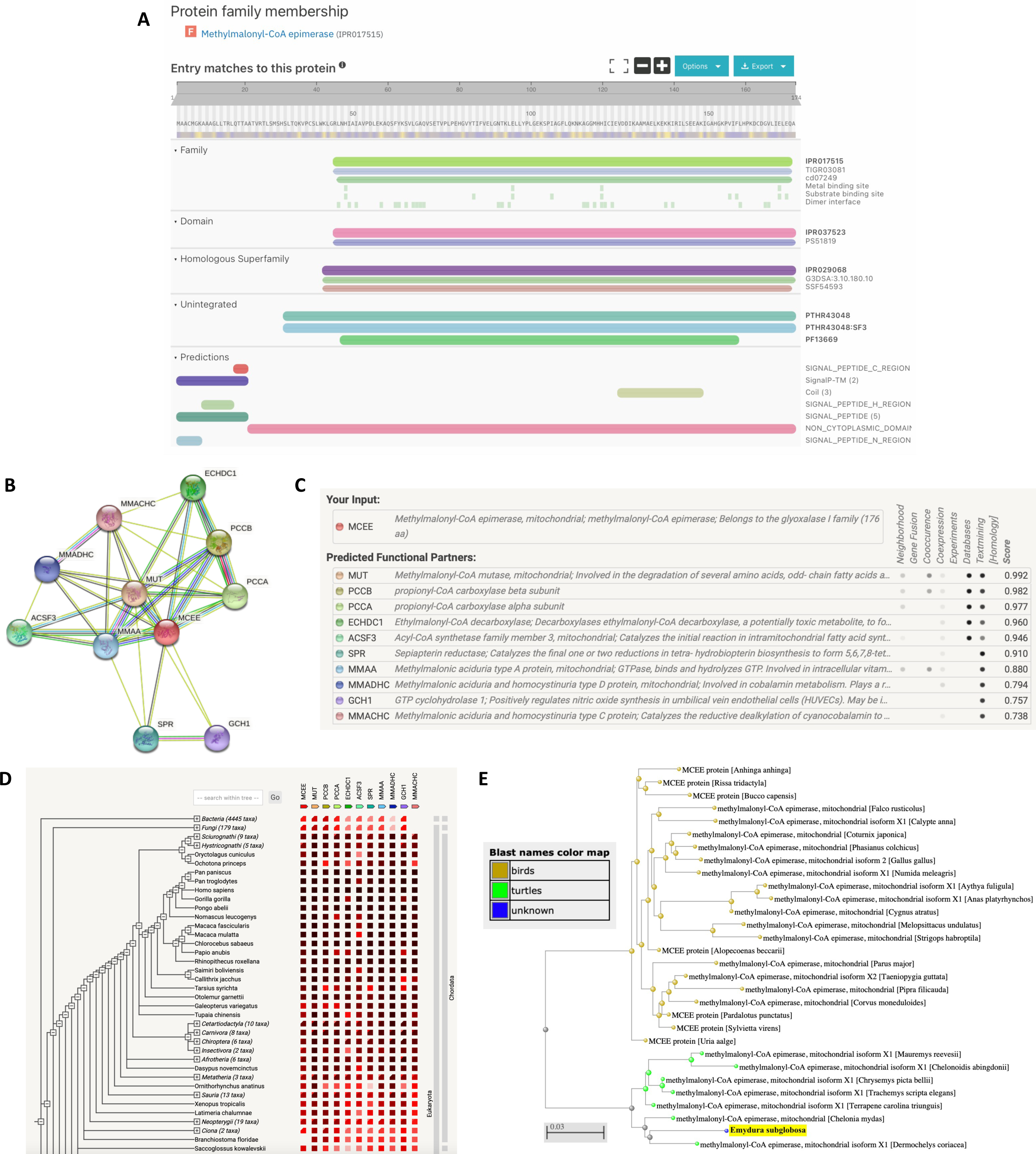
Functional analysis of the MCEE enzyme. (A) InterPro functional analysis of the enzyme. (B) STRING network of predicted protein-protein interactions in *Homo sapiens*. (C) List of functional partners predicted by STRING corresponding to B. (D) Gene co-occurrence of the enzyme. (E) BLAST phylogenetic tree built based on pairwise alignment.

To further analyze function, we also determined protein-protein interactions through the STRING database. The protein was predicted to be MCEE, which belongs to the glyoxalase I family (Figs 14B and 14C); this agreed with the InterPro protein family results (Fig 14A). The enzyme’s predicted functional partners included MUT (methylmalonyl-CoA mutase involved in degrading amino acids), MMAA (hydrolyzes GTP), ECHDC1 (decarboxylase), and several others (Figs 14B and 14C). These network links suggest the co-evolution of the non-homologous proteins since they are functionally related and could warrant additional analysis. Further research can also be done to determine whether they act in a complex or directly bind each other. Additionally, the gene co-occurrence suggests that these gene families have strong homology across the chordata phylum (Fig 14D). Finally, based on phylogenetic tree analysis of the methylmalonyl-CoA epimerase, *E. subglobosa* shared the same root as other turtle species, suggesting that these species came from a common ancestor (Fig 14E and S5 Fig).

We also wanted to analyze the subcellular localization of the protein based on any TMHs, signal sequences/peptides, and transit peptides. In WoLF-PSORT, a majority of the highest matches predicted the mitochondria, suggesting that this protein is likely located in the mitochondria (Fig 15A). There were no TMHs predicted and only non-cytoplasmic domains, so the protein does not localize to the plasma membrane (Fig 15B). The protein was predicted to not have a signal peptide sequence in both SignalP (Fig 15C) and TargetP (Fig 15E) but had very high probability for a mitochondrial transfer peptide (P = 0.9816, not shown). However, Phobius predictions did not agree with SignalP and showed a signal peptide from residues 1-20, so further validation and ribonucleic acid sequencing (RNA-seq) may be needed to gain confidence in whether this has a signal peptide (Fig 15D). Based on our analyses, the protein likely localizes to the mitochondria.

**Figure 15.**
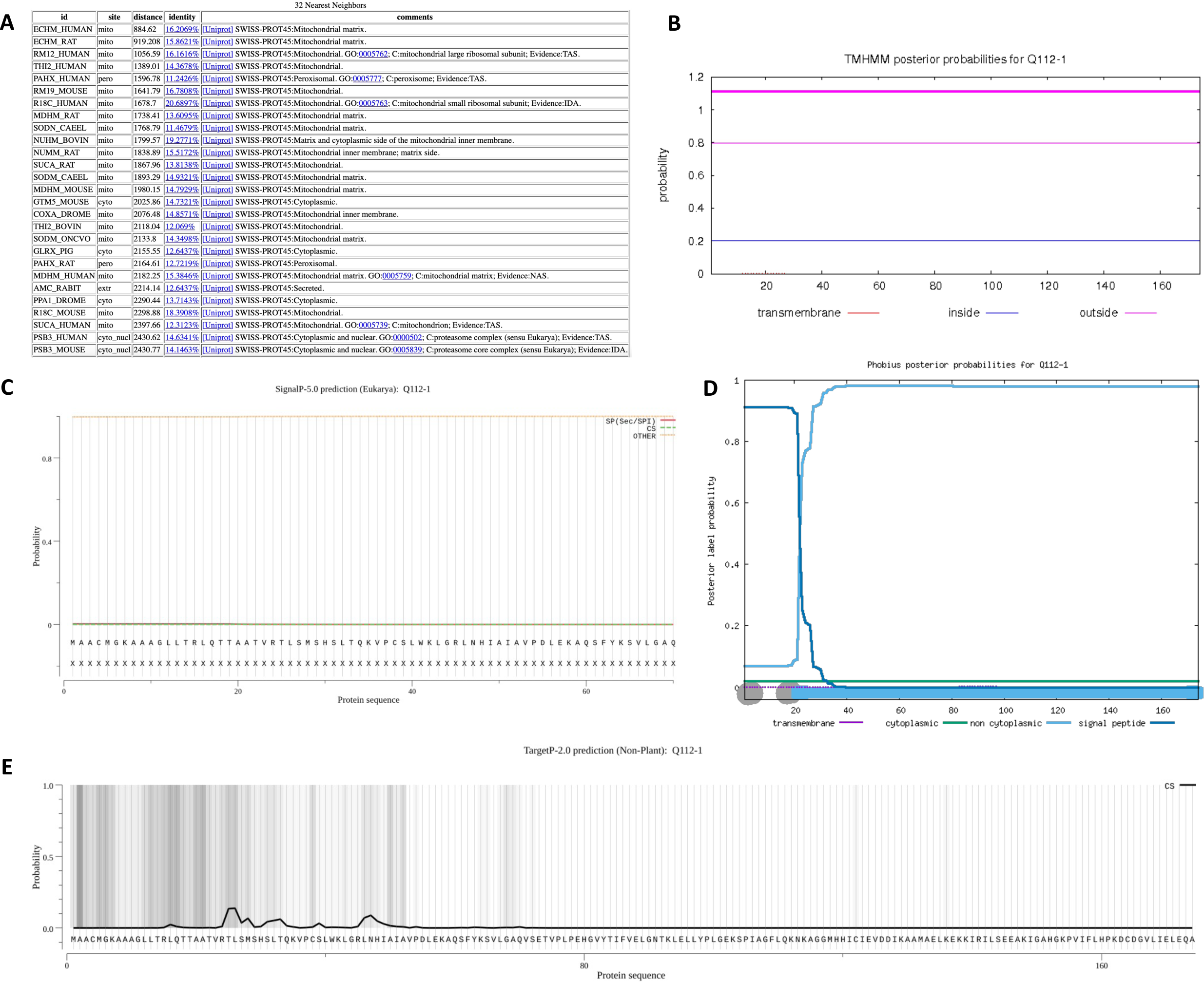
Subcellular localization of the MCEE enzyme. (A) WoLF-PSORT prediction of the protein localization site based on 32 nearest neighbors. Localization scores are ranked from the highest scoring matches (top) to lowest (bottom). (B) TMHMM prediction of TMHs. X-axis represents the amino acid number, and y-axis represents the probability that the amino acid is located within, outside, or inside the membrane. Probabilities >0.75 are significant. (C) SignalP analysis of signal sequences existing in the amino acid sequence of the polypeptide. (D) Phobius predictions of TMHs and signal peptides. X-axis represents the amino acid number, and y-axis represents the probability that the amino acid is transmembrane, cytoplasmic, non-cytoplasmic, and/or a signal peptide. Probabilities >0.75 are significant. (E) TargetP-2.0 prediction of N-terminal pre-sequences, signal peptides, and transit peptides.

After identifying the function and localization of the enzyme, we also wanted to understand its structure since protein structure heavily influences protein function. The SWISS-MODEL predicted a homo-dimeric structure that was highly conserved based on homology (Figs 16A and 16B). To validate the model, we examined the global and local quality estimates. The GMQE was 0.59, QMEAN was 1.44, and sequence identity was 81.06% (Fig 16B). Additionally, the local quality estimates had high probabilities in a majority of the regions; areas that dipped below 0.6 may have had poor resolution or were less conserved (Fig 16C). Notably, the homodimers were also extremely similar. Comparing the model to the non-redundant set of PDB structures, the QMEAN was within the average QMEAN scores for proteins of similar size (Fig 16D). The Ramachandran Plot also showed high favorability of the residues having a specific orientation, as shown by a majority of dots (97.31%) located in the dark green regions (Figs 16E and 16F). The MolProbity Score was near 0; the clash score, outliers, deviations, and bad angles were low (Fig 16F). PSIPRED predicted both alpha-helixes and beta-strands in the secondary structure, which was consistent with SWISS-MODEL (Fig 16G). Together, these data suggest a high-quality structure that could be used to model the methylmalonyl-CoA epimerase enzyme.

**Figure 16.**
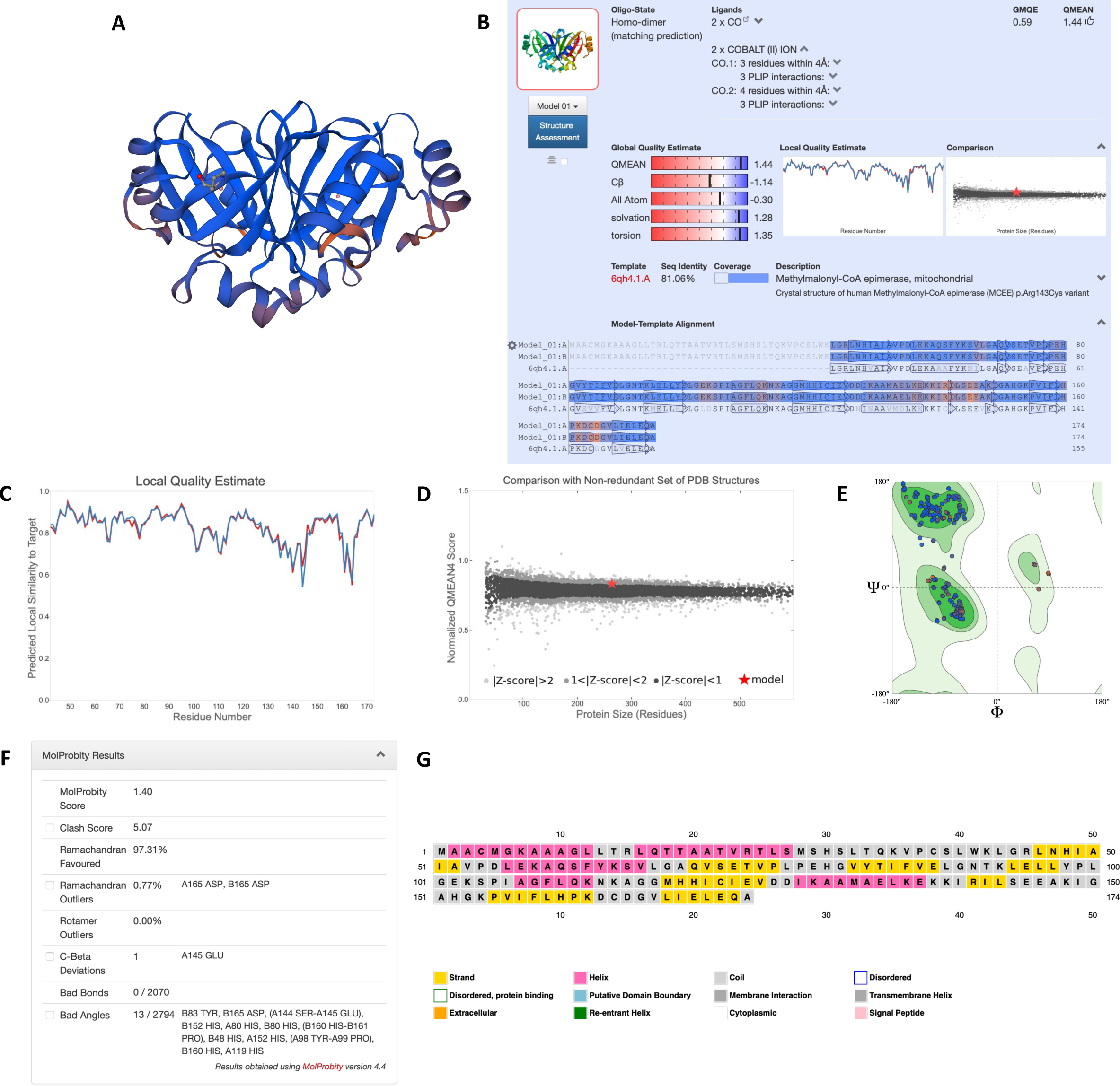
Homology modeling and structural predictions of the MCEE enzyme. (A) Three-dimensional homology model built by SWISS-MODEL. Blue regions are highly conserved, while orange regions are less conserved. (B) Oligo-state, ligands, global quality estimates, template, sequence identity, and coverage outputted by SWISS-MODEL. (C) Local quality estimate showing pair residue estimates. Similarities >0.6 are high-quality models. (D) Comparison with non-redundant set of PDB structures showing QMEAN scores for experimental structures that have been deposited of similar size. The red star is our model. (E) Ramachandran plot showing the probability of a residue having a specific orientation. Dots in the dark green regions represents high probability and a high-quality model. (F) MolProbity results to validate the model. A MolProbity Score close to 0 represents the resolution that one would see a structure of this quality. Clash score represents overlapping residues; a lower value is favored. Outliers represent values that extend outside the standard deviation; low values are also favored. Low values for bad bonds and angles are also favored. (G) Secondary structure prediction through PSIPRED.

## Discussion

Our study manually annotated and improved upon four predicted gene models in the ML679947.1 scaffold of the *Emydura subglobosa* genome. There is strong evidence that these genes encode a mitochondrial cholesterol side-chain cleavage enzyme, receptor for retinol uptake STRA6, lysyl oxidase homolog 1, and mitochondrial methylmalonyl-CoA epimerase--three of which fall under the enzyme class of oxidoreductases. Initial BLAST pairwise alignments and COBALT multiple sequence alignments showed these genes were highly conserved across homologous sequences, including those from turtles and other vertebrate species.

For the g19.t1 gene (CYP11A1), we validated the gene model predicted by AUGUSTUS and concluded that it was the most refined consensus model. We identified that the CYP11A1 gene encodes a mitochondrial cholesterol side-chain cleavage enzyme, as predicted by BLAST and InterPro, and is important in C21-steroid hormone biosynthesis. Previous studies have shown that the cholesterol side-chain cleavage enzyme catalyzes the initial, rate-limiting step of steroidogenesis by cleaving cholesterol and generating pregnenolone--a precursor for essential steroids in the adrenal, gonads, and placenta such as progesterone, cortisol, and aldosterone [28, 29]. In addition to steroid production, the cholesterol side-chain cleavage enzyme also has been demonstrated to play a role in the morphology of the mitochondria, particularly its cristae shape which is important for oxidative phosphorylation [28]. This study agreed with our WoLF-PSORT (Fig 3A) and TargetP (Fig 3E) results predicting a mitochondrial subcellular localization. Additionally, mutations in the CYP11A1 gene can lead to life-threatening adrenal and gonadal insufficiencies, disruptions in the corticosterone circadian rhythm, and blunted stress response [29–31]. These findings highlight the evolutionary importance of the CYP11A1 gene through its highly conserved sequence.

We also manually annotated and improved upon the g24.t1 gene (STRA6) by removing an exon containing 17 extra residues when comparing to homologous sequences. The STRA6 gene was characterized to encode a receptor for retinol (Vitamin A_1_) uptake, as predicted by BLAST and InterPro, and plays a role in retinol transmembrane transporter and signaling transporter activity. Not only is receptor STRA6 revealed to be a transmembrane pore mediating the bidirectional transport of retinol, but it also is a receptor activated by the retinol binding protein complex and induces downstream activation of STAT target genes [32, 33]. Studies have conveyed the importance of retinol in vision, anti-aging, cell morphogenesis, differentiation, proliferation, and cancer prevention [34, 35]. Mutations in STRA6 have been shown to result in dysfunction of vitamin A transport, type 2 diabetes, and Matthew Wood Syndrome [32]. Additionally, previous structural analyses characterized STRA6 as containing one intramembrane and nine transmembrane helices in a dimeric structure [36], which corroborates with our results showing 10 TMHs. However, further analysis using a digestion assay could elucidate which helix corresponds to the intramembrane helix in the STRA6 gene. Also, our SWISS-MODEL results did not agree with [36], which was expected because we had very little confidence in the model. Through the elucidation of the STRA6 gene, we can understand the potential anti-aging, anti-cancer, nutritive effects of retinol uptake in turtles.

Following manual annotation and generating a refined consensus model, we revealed that the g32.t1 gene likely encoded a lysyl oxidase homolog 1 (LOXL1) protein. Our InterPro results predicted a conserved region, our GO terms predicted copper ion binding, and our localization analyses predicted a signal peptide. These three findings corroborate with a previous study showing that the LOX family genes have a conserved catalytic C-terminal domain, conserved copper-ion binding domain, and signal peptide across all LOX proteins [37]. A study elucidated the protein-protein interactions between LOXL1 with the other LOX family members as well as proteins involved in elastic fiber (MFAP2/5 and FBLN5) and collagen formation (BMP1, PCOLCE), which supported our STRING results [38]. Furthermore, two studies have shown that LOXL1 directly interacts with Fibulin-5 (FBLN5), and BMP1 cleaves Pro-LOXL1 to form LOXL1, which also corroborates with STRING [39, 40]. LOXL1 is a highly important enzyme in elastic fiber biogenesis in the extracellular matrix, crosslinking of collagen fibrils, and matrix remodeling during injury or tumorigenesis [40]. Knockdown of LOXL1 has been demonstrated to exert oncogenic roles and can lead to exfoliation syndrome and glaucoma in humans [41, 42]. Thus, the highly conserved LOXL1 gene plays a significant role in tumor suppression, wound healing, and vision not only in turtles but also in many other vertebrates (as shown by our phylogenetic analysis).

Finally, through our analysis of the g112.t1 gene (MCEE), we found that it likely encodes methylmalonyl-CoA epimerase, an essential enzyme involved in branched-chain amino acid and odd-numbered fatty acid catabolism by converting (2R)-methylmalonyl-CoA to (2S)-methylmalonyl-CoA [43]. This pathway eventually leads to the downstream formation succinyl-CoA for the citric acid cycle. A previous paper studied the crystal structure of this enzyme and found that MCEE exists in a stable dimer with an 8-stranded beta sheet consisting of monomers folded into two tandem beta-alpha-beta-beta-beta modules [43]. This was very consistent with the SWISS-MODEL homology model. MCEE deficiency has been shown to lead to methylmalonic aciduria and central nervous system damage in humans [43, 44]. Therefore, MCEE is an important gene for catabolism, so turtles can respond and adapt to their changing needs for energy.

Elucidating the *E. subglobosa* genome through manual annotation contributes to the increasingly important field of conservation genomics and biodiversity [45]. With over half of the turtle species threatened by extinction, we characterized four highly conserved genes to further contextualize the reproductive biology of turtles and document their genetic diversity. The genes presented in this study contribute to the longevity phenotype of turtles and can be used by geneticists to direct conservation efforts towards critically endangered species. Furthermore, knowledge of the genes that are involved in the reproductive success and longevity of turtles can be further applied to clinical settings such as disease treatment in humans. Overall, our genomic analysis of the *Emydura subglobosa* genome advances management and conservation efforts aimed to save currently endangered turtle and vertebrate species, provides clinical importance and application for human therapies, and adds significant progress towards a fully annotated genome.

### Limitations & Future Directions

Through our study, we were limited to only computational and homology-based strategies. Another limitation was that we only analyzed one scaffold of the genome. We propose the use of RNA-seq experiments to further contextualize the *E. subglobosa* genome and contribute to our expanding knowledge about turtle biodiversity. Additionally, we could also manually annotate and refine different genes located in other scaffolds or other turtle species.

## Acknowledgements

Thank you to Professor Matteo Pellegrini, Leroy Bondhus, and Noah Alexander for their teaching and guidance throughout MCD BIO 187AL.

## Supporting information

S1 Fig. COBALT phylogenetic tree of the mitochondrial cholesterol side-chain cleavage enzyme. The COBALT multiple alignment tree is comparable to the BLAST pairwise alignment tree in Fig 2F. Both demonstrate the different groupings of species such as salamanders, snakes, lizards, birds, vertebrates, and turtles. Both show that the E. subglobosa are an outgroup of the turtles.

S2 Fig. COBALT phylogenetic tree of the receptor for retinol uptake STRA6. The COBALT multiple alignment tree is comparable to the BLAST pairwise alignment tree in Fig 6F. Both demonstrate the different groupings of species such as vertebrates, birds, and turtles. Both show that the E. subglobosa are an outgroup of the turtles.

S3 Fig. COBALT phylogenetic tree of lysyl oxidase homolog 1. The COBALT multiple alignment tree is comparable to the BLAST pairwise alignment tree in Fig 10F. Both demonstrate the different groupings of species such as caecilians, marsupials, whales/dolphins, bats, carnivores, rodents, placentals, monotremes, lizards, snakes, birds, and turtles. Both show that the E. subglobosa are an outgroup of the turtles.

S4 Fig. AmiGO 2 gene ontology graphs for the mitochondrial MCEE enzyme. (A) Biological process L-methylmalonyl-CoA metabolic (GO:0046491). (B) Molecular function methylmalonyl-CoA epimerase (GO:0004493).

S5 Fig. COBALT phylogenetic tree of the mitochondrial MCEE enzyme. The COBALT multiple alignment tree is comparable to the BLAST pairwise alignment tree in Fig 14E. Both demonstrate the different groupings of species such as birds and turtles. Both show that the E. subglobosa share the same root as the turtles.

## Notes

### Competing Interest Statement

The authors have declared no competing interest.

