## Supplementary figures and images for "Elucidating the genome of *Emydura subglobosa* through manual annotation and computational biology tools"

### S1 Fig

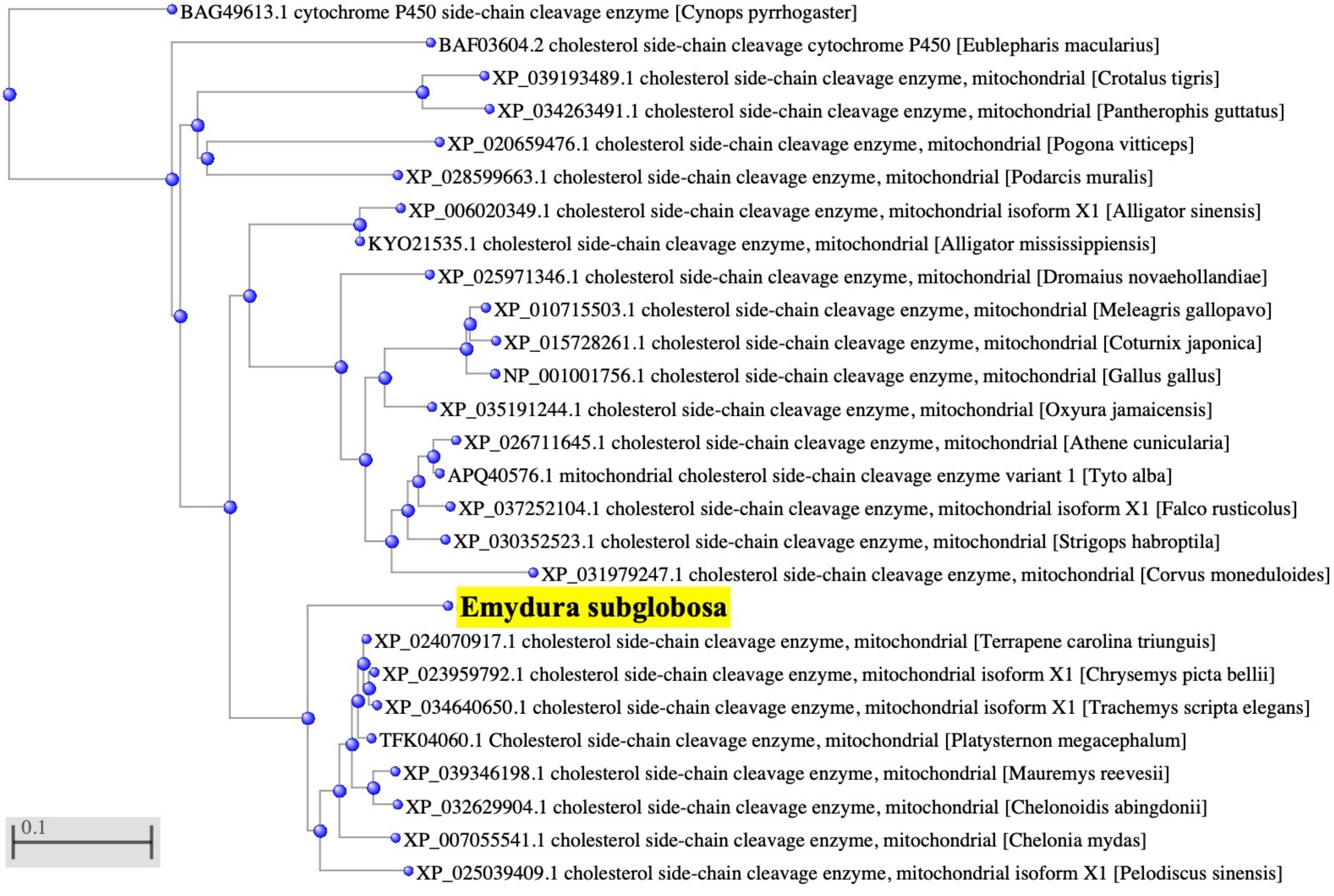

### S2 Fig

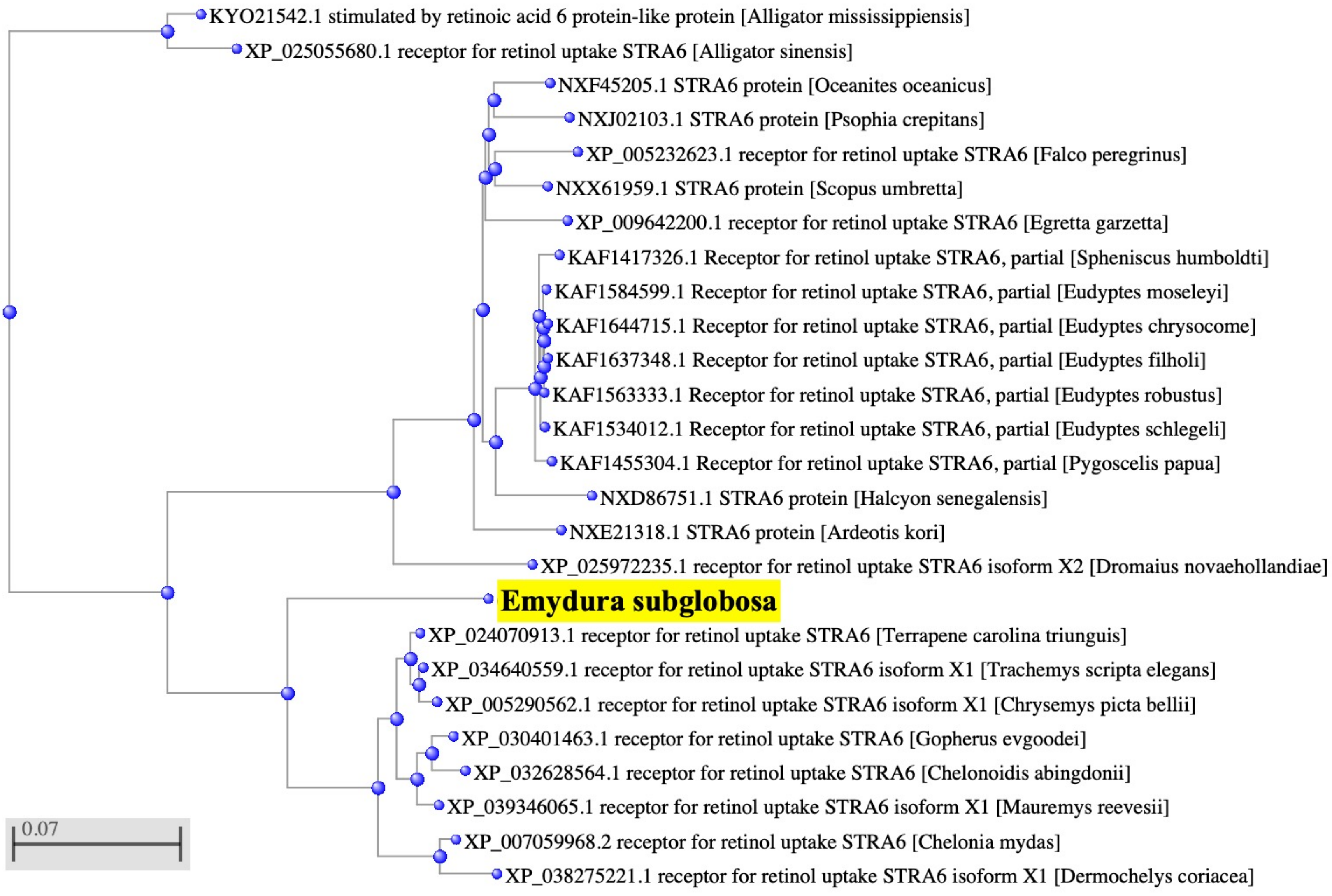

### S3 Fig

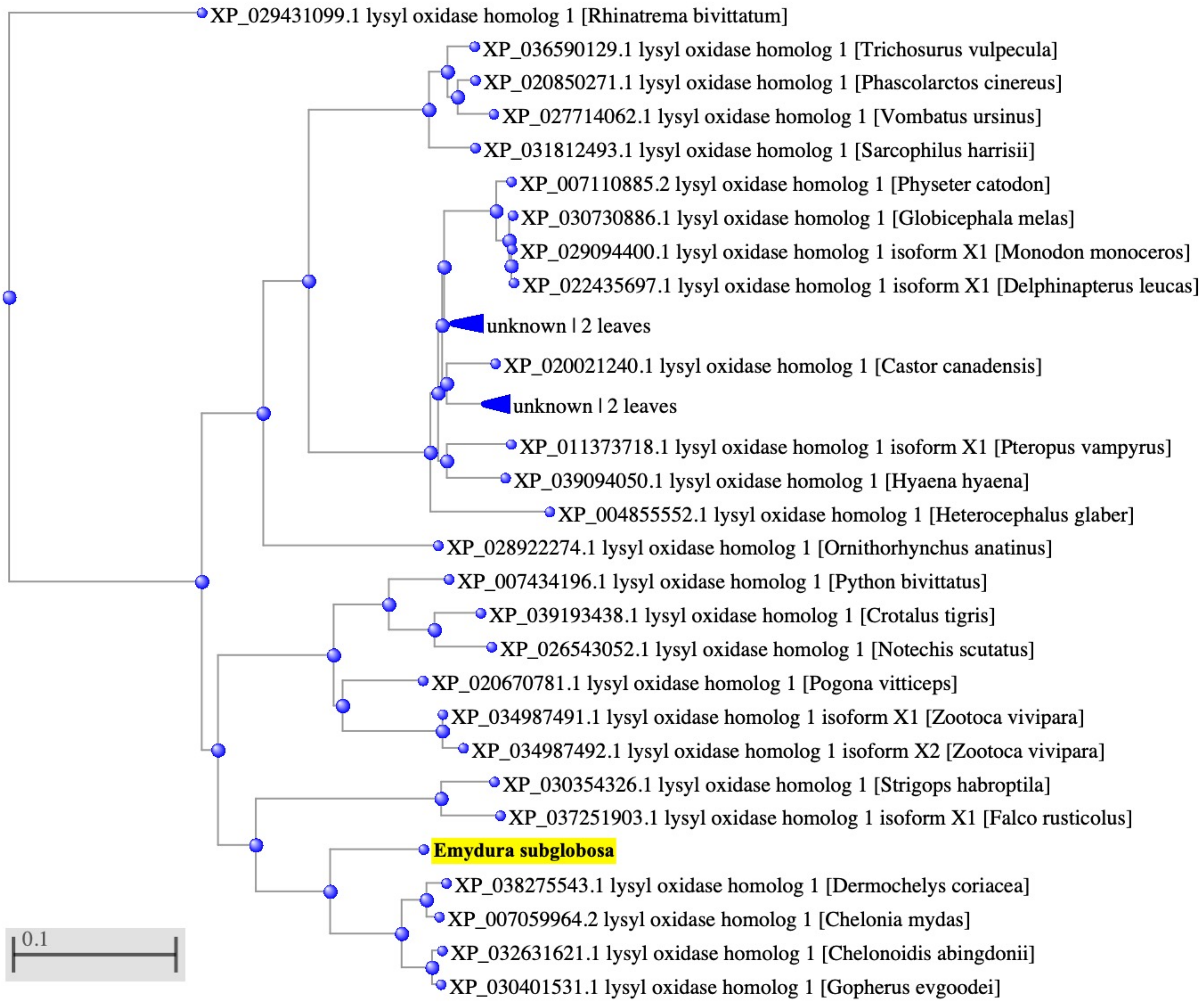

### S4 Fig

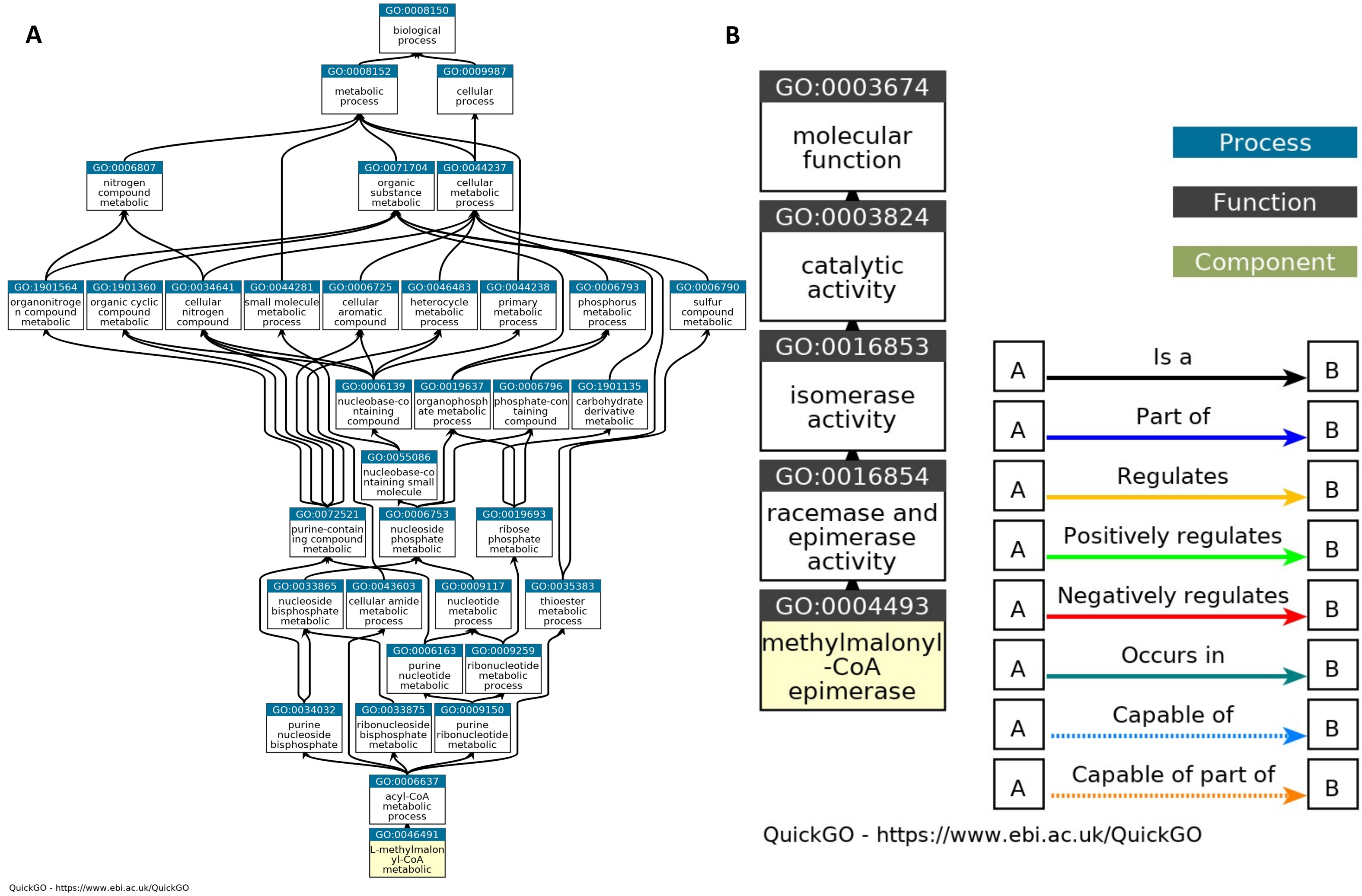

### S5 Fig

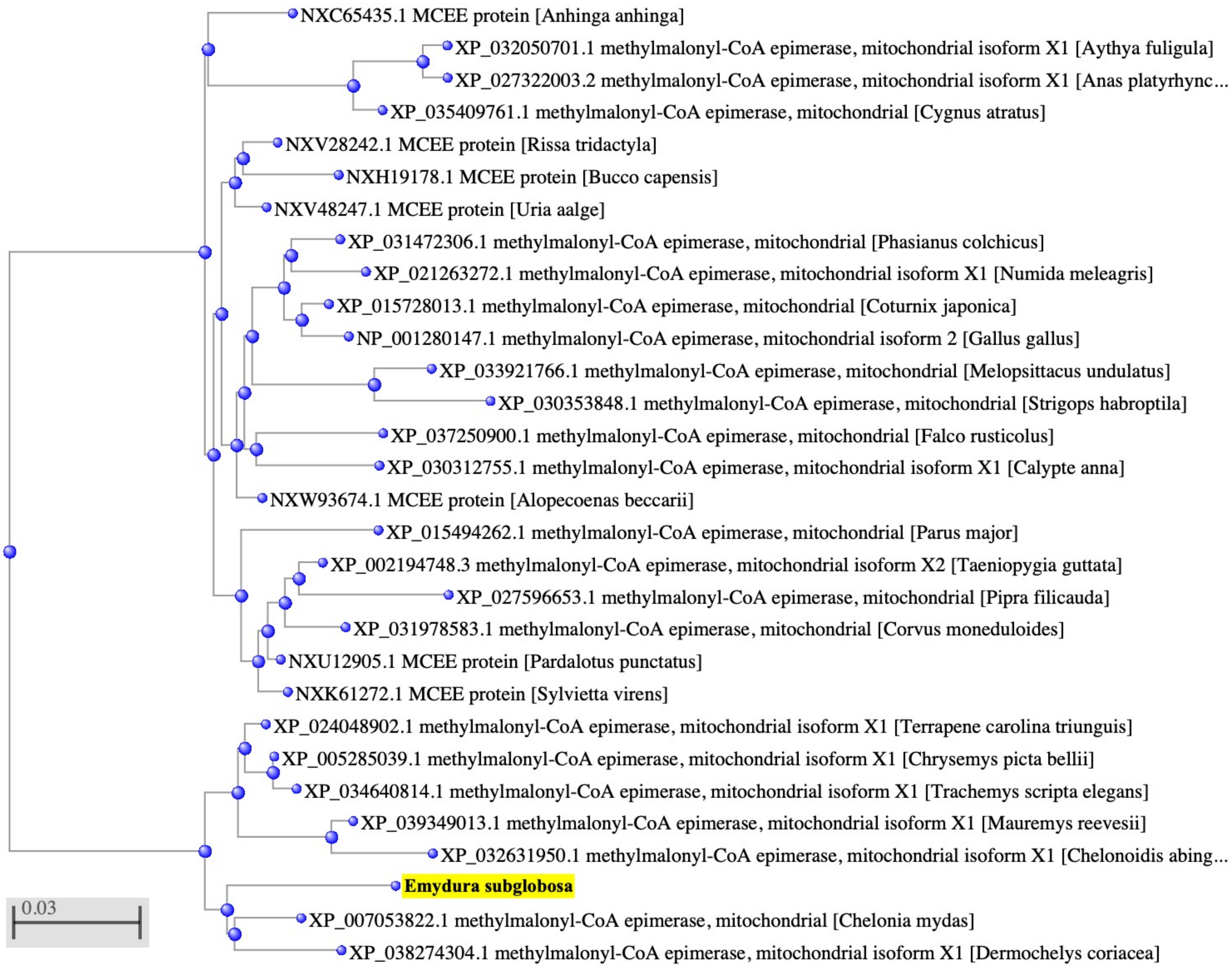
